# MorphoNet 2.0: An innovative approach for qualitative assessment and segmentation curation of large-scale 3D time-lapse imaging datasets

**DOI:** 10.1101/2025.02.21.639560

**Authors:** Benjamin Gallean, Tao Laurent, Kilian Biasuz, Ange Clement, Noura Faraj, Patrick Lemaire, Emmanuel Faure

**Author notes:** These authors contributed equally to this work.

## Abstract

Thanks to recent promising advances in AI, automated segmentation of imaging datasets has made significant strides. However, the evaluation and curation of 3D and 3D+t datasets remain extremely challenging and highly resource-intensive. We present MorphoNet 2.0, a major conceptual and technical evolution designed to facilitate the segmentation, self-evaluation, and correction of 3D images. The application is accessible to non-programming biologists through user-friendly graphical interfaces and works on all major operating systems. We showcase its power in enhancing segmentation accuracy and boosting interpretability across five previously published segmented datasets. This new approach is crucial for producing ground-truth datasets of discovery-level scientific quality, critical for training and benchmarking advanced AI-driven segmentation tools, as well as for competitive challenges.

## Introduction

Recent advances in optical time-lapse microscopy now enable the 3D capture of life’s temporal dynamics, from molecular assemblies to entire organisms^1^, in cells, embryos, and aggregates. These technologies have thus become crucial for cell and developmental biology. The automated segmentation and tracking of these complex datasets is, however, necessary for their interpretation and quantification^1^.

Recent progress in the 3D segmentation of animal and plant cells^2^, or organelles^3^ has made this process highly efficient, despite the persistence of algorithm-dependent systematic errors^4^. The processing of time-lapse datasets, however, remains challenging. The accumulation over hundreds of time points of even few residual segmentation errors can strongly affect data interpretation and notably disrupt long-term cell tracking. Following the excitement generated by this new generation of imaging systems, which offers the ability to image living systems at unprecedented spatial and temporal scales, a glass ceiling has now been reached due to the quality of these reconstructed data, which fails to scale to real research exploitations.

Beyond the reconstruction requirements of individual research projects, proper metrics are essential for evaluating the performance of bioimage analysis algorithms^5^. However, the training and benchmarking of next-generation AI-based segmentation tools fundamentally depend on the availability of accurate ground truth data^6^. The critical bottleneck is the lack of a massive amount of 3D high-quality reconstructed data to train these systems.

We have decided to take on this challenge and develop a tool to produce high-precision and high-quality 3D datasets for AI. Two major barriers must be overcome to reach this milestone. The first concerns the **evaluation** of dataset reconstructions: how can we measure the accuracy of segmentations obtained by automated algorithms for these complex 3D and 3D+t datasets? The second focuses on the **curation** of reconstructed data: how can we correct and significantly enhance segmentations to achieve reliable reconstructions that are essential for major scientific breakthroughs?

2D imaging has long been a cornerstone of biological research, driving significant scientific discoveries with visual validation being largely adequate, as it could be performed directly on a 2D screen. The traditional method involves visually comparing the reconstruction by overlaying it with the acquired data, as seen in classic software like ImageJ^7^, Napari^8^ or Ilastik^9^. When discrepancies arise, experts can perform curation and manually adjust the pixel values to align with the desired class (e.g., a cell).

In the context of 3D and 3D+t data, the technique encounters significant obstacles that hinder reliable scientific analysis. The 3D structure of images, inherently unsuited to the 2D digital environment, demands numerous compromises that restrict their interpretability. Projecting 3D objects onto a 2D screen causes data masking, requiring experts to perform extensive manipulations to verify accuracy. Additionally, the high number and density of 3D objects further complicate the process, as visual interference from overlapping objects increases during these adjustments. As a result, voxel-level (3D pixel) curation becomes almost unfeasible using conventional methods, compelling experts to revert to working in 2D. This approach becomes very time-consuming and therefore does not allow to obtain a satisfactory reconstruction quality for scientific use. A final obstacle is caused by the much larger size of 3D datasets, which significantly lengthens processing time for each curation action. This computational burden becomes daunting for 3D + t time-lapse datasets, as error propagation between consecutive time points rapidly increases the number of segmentation and object tracking errors needing correction. Restricting the application of advanced image processing tasks to subsets of the image may alleviate this issue.

An unsupervised objective assessment can be derived from an *a priori* knowledge of expected spatial (e.g. smoothness of contours, shape regularity, volume…) or temporal (e.g. stability and smooth evolution, lifetime of objects…) features of the objects^10^. Features of the object boundaries (e.g. contrast between inside and outside of an object) can also be computed^10^. More sophisticated strategies have been proposed when no statistically-relevant a priori knowledge can be drawn^11^. Since curation is typically carried out by expert biologists with limited programming skills, the computation of image features and the projection of relevant metrics onto individual segmented objects should be accessible through user-friendly interfaces. An alternative approach for interacting with 3D segmented images is therefore necessary.

## Results

We previously developed a novel concept of morphogenetic web browser, MorphoNet^12^, to **visualize** and **interact** with 3D+t segmented datasets through their meshed representations. This user-friendly web solution required no installation and was suited for datasets of moderate size (up to a thousand cells over a few hundred time points). Since its release, this platform has been used in a variety of morphological studies in plant and animal systems^13,14^ and has been instrumental to interpret the relationship between gene expression and complex evolving cell shapes^15–17^. This web version was restricted to visualizing and interacting with segmented datasets through their precomputed 3D dual meshes. This approach offered significant benefits in terms of online 3D rendering efficiency and data management but it also came with limitations. For example meshes are simplified representations of the segmented object that may not capture fine details of the original volumetric data and that cannot be validated without comparison to the original raw intensity images. Meshed representations can also make it challenging to perform the precise computations required for detailed geometric and topological analyses^18^. Exploration of larger datasets was nonetheless constrained by the computational resource limitations of internet browsers.

We present here MorphoNet 2.0, a major conceptual evolution of the platform, designed to offer powerful tools for the **reconstruction**, **evaluation**, and **curation** of 3D and 3D+t datasets. This innovative approach leverages the richness and redundancy of information embedded within these complex datasets. It addresses the two previously identified challenges: **evaluating** 3D segmented data without relying on manual ground truth by fully harnessing the data’s richness, and enabling semi-automated **curation** of 3D data, now achievable with just a few clicks. This enables us to achieve data quality at a level suitable for scientific discovery. We showcase the impact of these advancements by revisiting and enhancing five published animal and plant datasets previously regarded as ground truth^6^.

We feature a new standalone application running on all major operating systems that exploits the resources of the user’s local machine to explore, segment, assess and manually curate very large 3D and 3D+t datasets. Like the web version, the MorphoNet application uses the power and versatility of the Unity game engine and includes its main features^12^. By overcoming web-based limitations, the application can handle more complex datasets including heavier segmented voxel images and up to several tens of thousands of objects on a research laptop (Fig. 1h, and Supp. Table 1). The standalone application allows users to directly explore private data stored on their own computer, without need for an upload to the MorphoNet server. Subsequent data sharing with other researchers or a wider public in an open science process is facilitated by the MorphoNet server upload functions of the standalone.

**Figure 1:**
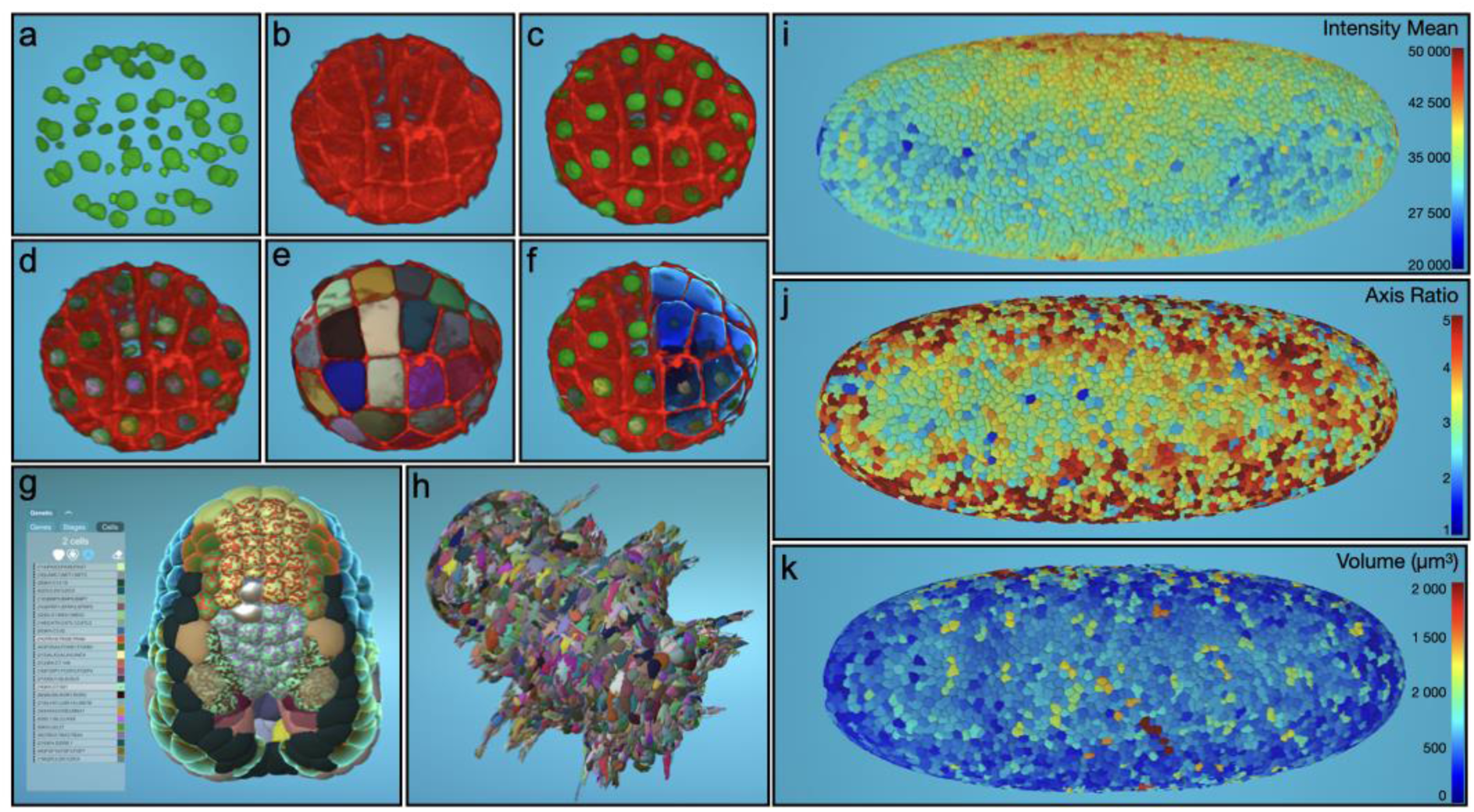
Illustration of the visualization of datasets of various complexity and nature in the MorphoNet standalone application. *a-f. Visualization of a 64-cell stage Phallusia mammillata embryo with labeled cell nuclei and cell membranes. a-c intensity images showing the nuclei (a), the membranes (b) or both (c). d. same as c. with additional nuclei segmentation obtained with the Binarize plugin (see Supp. Mat. for the full description of curation plugins). e. same as b. With additional membrane segmentation obtained with the Cellpose 3D plugin. f. same as c. with a combination of several rendering possibilities of cell and nuclei segmentations*. *g. Multi-colored shaders allow the simultaneous visualization of the expression patterns of multiple genes extracted from the ANISEED^17^ database and of tissue fate information. Ascidian embryo^16^ at stage 15 (mid neurula); cells with a single color are coloured with larval tissue fate; multi-coloured cells are coloured with both larval tissue fate and the expression of selected genes*. *h. Visualization of a six days post-fertilization Platynereis dumerilii embryo^20^ imaged by whole-body serial block face scanning electron microscopy followed by the automated whole-cell segmentation of 16000 cells*. *i-k. Visualization of a cell cycle 14 Drosophila melanogaster embryo imaged with SiMView microscopy and segmented with RACE^21^. i. Projection on each segmented cell of the mean image intensity. j. Projection on each segmented cell of the ratio between the length of the major and the minor axes of the ellipse that has the same normalized second central moments as the segmented cell. k. projection of the cell volume*.

MorphoNet is designed with a strong emphasis on user-friendliness, making its use highly intuitive for experimental biologists with no coding experience (See Supp. Movies). Every feature and interface element has been crafted to ensure a smooth, straightforward user experience, allowing both beginners and experts to utilize its capabilities efficiently. Additionally, the solution is fully open-source allowing bio-image analysis to develop their own features (See Supp. Table 2).

One of the key features of this new standalone application is the automatic integration of a duality between the 3D segmented images and their corresponding meshes (Fig 2.). This feature is achieved by linking Python, dedicated to high-performance image processing, with the Unity Game Engine, fully optimized for seamless interaction with the dual meshes. This integration enables access to state-of-the-art reconstruction^2,19^ tools, including those leveraging AI libraries, while also addressing the challenges of interacting with 3D images by harnessing the powerful interactive capabilities of game engines.

**Figure 2:**
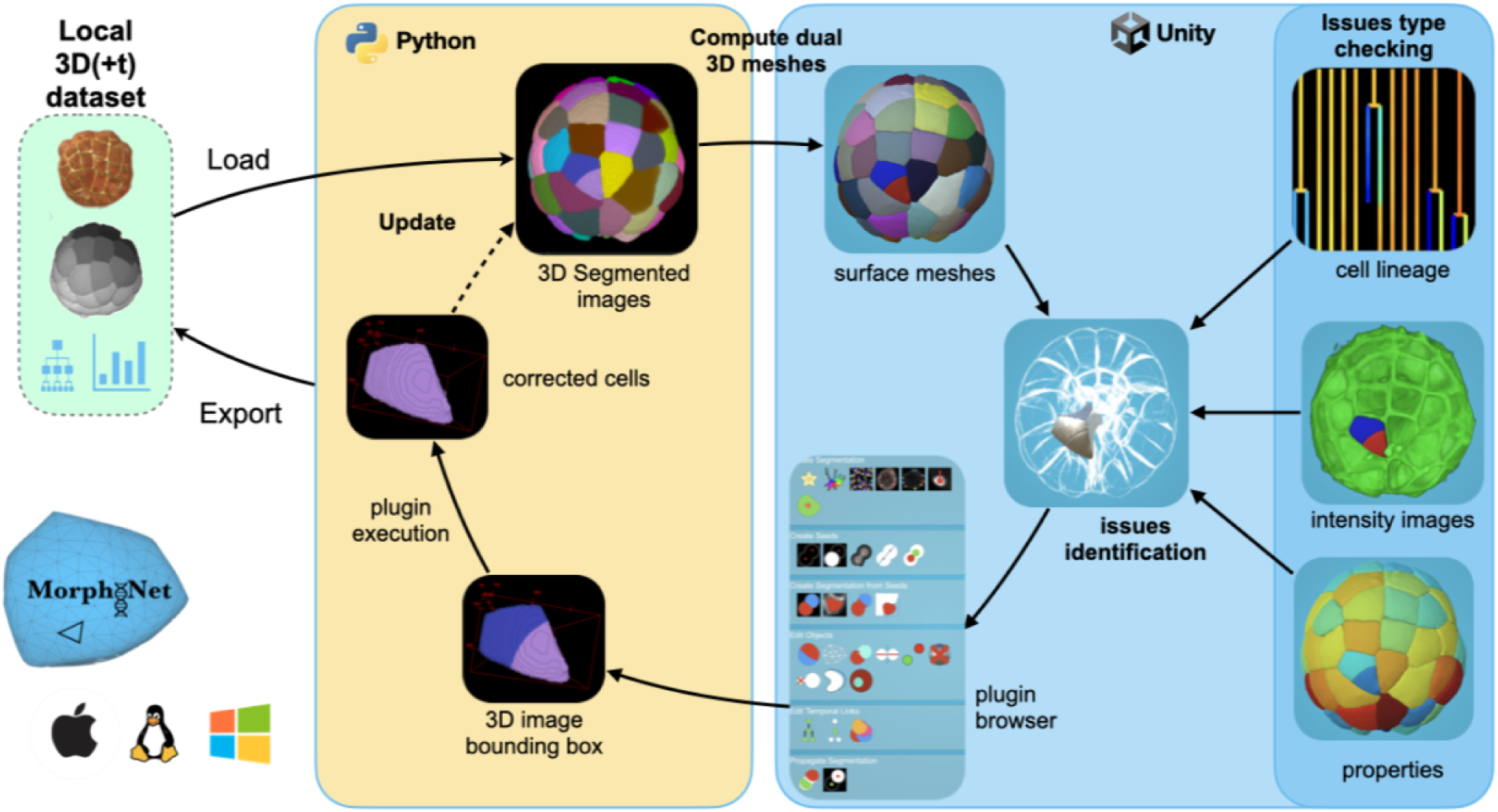
MorphoNet Standalone Schema: From the local data (loaded from the green box), the MorphoNet Standalone application computes first the dual meshes for each of the segmented objects (in the module python in the yellow box). Then, using the 3D viewer (in the blue box), users identify detection, segmentation or tracking issues using, if necessary, the cell lineage information, the raw images and/or properties computed on the segmented objects. Errors are then corrected by choosing and executing the appropriate image processing plugin from the curation menu. Finally, new meshes are computed from the result of the plugin execution to update the visualization.

To ensure users can consistently assess segmentation quality, raw intensity images are also available by superimposition. This provides access to the entire workflow, from raw image acquisition to meshed segmentation, allowing simultaneous exploration and visualization of both meshed and voxel images (Fig. 1a-f).

We also leveraged Unity’s scene management tools to implement simultaneous visualization of multiple scenes. This feature enables the display of interactive cell lineages in a dedicated, fully connected window, offering essential insights into cellular trajectories within 3D+t datasets.

### Automatic error detection

Efficient dataset curation requires the rapid identification of segmentation errors in large 3D or time-lapse datasets containing thousands of segmented objects. MorphoNet 2.0 tackles this critical challenge by introducing unsupervised metrics for objective, automated assessment, eliminating the subjectivity, inefficiency, and inconsistency inherent in manual visual inspections. These metrics leverage prior knowledge of generic data properties - such as shape regularity, contour smoothness, and temporal stability of shapes and volumes - to deliver scalable and consistent analysis. Beyond improving curation efficiency, these metrics offer quantitative scores that enable systematic comparisons across datasets, experiments, and tools. Integrated into MorphoNet, they support benchmarking, algorithm refinement, and reproducibility. Using the scikit-image library^22^, we automatically compute object properties from both segmented and intensity images. These properties quickly help identify outliers, which often correspond to segmentation errors. While the traditional approach uses some metrics to evaluate global distributions, it is crucial for curation purposes to apply these properties at the level of each individual segmented object. Thus, these properties, including shape metrics like volume, convexity, and elongation, and intensity metrics such as mean voxel intensity within or around segmented objects, can be easily projected onto meshed objects for visualization (Fig. 1i-k).

Segmentation quality assessment is further enhanced by calculating three categories of properties for each dataset: 1) Morphological features – including volume, convexity, roughness, and elongation, 2) Intensity-based measurements – within and around each object in the original acquisition images, such as mean intensity, homogeneity, or deviation at segmentation borders; 3) Temporal features – such as object lifetime or cell lineage distances^16^.

By projecting these values onto segmented object meshes, outliers are readily identified as prime candidates for curation. These values can also be exported and plotted to reveal distributions, providing an unsupervised assessment of overall segmentation quality. This process is exemplified in various use cases described below.

To evaluate the relevance of these unsupervised metrics, we compared their distributions in the published and curated datasets against a manually annotated gold standard in Use Case 1. The curated segmentations exhibited higher Intersection over Union (IoU) scores (Supp. Fig. 3). Although these unsupervised metrics are not flawless predictors of segmentation accuracy, they show a strong correlation with standard quality indicators such as IoU. This confirms their practical utility as reliable proxies for segmentation quality and valuable tools to guide efficient manual curation.

A key limitation of relying solely on unsupervised metrics is the risk of circular logic: corrections or model training based on masks deemed “good” by these metrics may promote homogeneity without improving biological accuracy. This can lead to segmentations optimized for metric conformity rather than real quality. To avoid this, MorphoNet’s metrics are intended as guidance tools for flagging candidate errors, which should then be validated through expert curation, ground truth comparisons, or biological plausibility.

### Biocuration

The execution time of algorithms in 3D image processing limits the optimization of their parameters. Additionally, the inherent heterogeneity within 3D images frequently prevents achieving consistently high-quality results across the entire dataset. To tackle these challenges, we defined in MorphoNet the notion of multi-type representation layers, able to combine and interoperate 3D image intensity layers with mesh-based layers. This dual representation enables seamless interaction with meshes, facilitating the rapid identification and precise selection of objects requiring curation. By focusing processing efforts on these selected regions rather than the entire image, MorphoNet significantly reduces the time and effort required, streamlining the curation workflow for complex 3D+t datasets.

The meshed versions of the segmentations are used for 3D rendering, for the identification and selection of objects needing curation, and to launch the needed curation algorithms, which are performed on the segmented images (Fig. 2). A significant acceleration of operations on 3D images is achieved by targeting the processing activities on a subpart of the image, for instance on the bounding box of an object or of a set of objects. This approach enables results to be generated in just seconds for most tasks, even with highly complex 3D data. Given the heterogeneity within 3D images, it is challenging to find a single set of parameters that works effectively across the entire image. This approach makes it easy to run image processing algorithms with different parameters in different subregions of the image.

Editions of objects necessitating curation can then proceed using dedicated plugins. The curation module of MorphoNet has an expandable open-source Python plugin architecture and provides user-friendly graphical interfaces accessible to experimental biologists with limited programming skills. To support advanced image processing techniques powered by deep learning, which have an increasing impact on image analysis, the default installation package includes training libraries like Scikit-learn, PyTorch, and TensorFlow. Plugins can take the raw intensity image into account to perform 3D image processing tasks as exemplified by the integration of popular segmentation tools, such as Stardist^19^ and Cellpose^2^. The full list of plugins can be found in the Supplementary Materials. Users and developers can expand the current plugin list by creating their own through an easy-to-use python template (See Supp. Table 2).

To help the user identify suitable plug-ins, we grouped them into seven categories. The first five are used to create or edit segmentations of 3D datasets, the last two are dedicated to the temporal propagation of corrections in 3D+t datasets. All categories will be exemplified in the five use cases below. Supp. Fig. 1. illustrates how the majority of segmentation and tracking errors can be corrected using plugins, single handedly or in combination. For example, an under-segmentation can be corrected by running locally the *Cellpose* plugin as in use case 2 or the Temporal Propagation plugin (*Prope*) as in use case 5.

1- *De novo Segmentation* plugins: these plugins create a *de novo* segmentation from the original intensity images (or from a specified region of interest of the intensity images). This category includes segmentation tools such as Cellpose 3D or Stardist as well as simpler intensity thresholding. As described below, MorphoNet plugins allow to specifically target the application of these tools to error-rich regions.

2- *De novo Seed* plugins: these plugins automatically create seeds (i.e., 3D points in the 3D images), which can then be used to initiate segmentation tasks. Additionally, seeds can also be added manually using the interface.

3- *Segmentation from Seeds* plugins: these plugins are used to perform local segmentation using predefined seeds, for example by using a watershed algorithm.

4- *Segmentation correction* plugins: these plugins are used to correct the main classes of segmentation errors on selected segmented objects. This includes the fusion of over-segmented cells, the resolution of small artifactual cells or the splitting of under-segmented cells.

5- *Shape transform* plugins: these plugins are used to edit the shape of selected segmented objects. This includes classical morphological operators (such as dilate, erode, etc.) or the manual deformation of a cell’s shape.

6- *Propagate Segmentation* plugins: these plugins are used to propagate an accurate segmentation at a specific time point to erroneous segmentations at later or earlier time points. Examples: Propagate eroded cell masks to correct under segmentation errors.

7- *Edit Temporal Links* plugins: these plugins are used to curate cell lineage trees by creating or editing temporal information. Examples: Create temporal links using the spatial overlap between homologous objects at successive time points.

### Use cases

To highlight the efficiency of MorphoNet plugins in detecting and correcting the main types of errors across a broad range of organisms, we present five examples from previously published fluorescent live imaging datasets, representing various levels of quality and complexity. These examples collectively showcase MorphoNet’s powerful segmentation quality assessment and error detection capabilities in different contexts, ranging from low-quality nuclear segmentations to high-quality whole-cell segmentations. While these examples frequently involve the use of Cellpose, the current state-of-the-art tool for 3D dataset segmentation, the focus is not on emphasizing the tool’s power but on demonstrating how advanced tools can be enhanced through integration into MorphoNet’s sophisticated graphical plugin interfaces.

The first use-case introduces two unsupervised metrics for assessing dataset quality in the absence of ground truth on a time-lapse light-sheet microscopy movie of a developing *Tribolium castaneum* embryo. Using these metrics, we demonstrate the power of an iterative segmentation and training strategy. By applying a Cellpose model, trained on a curated subpopulation, to the entire dataset, we improve segmentation quality.

The second use-case presents unsupervised whole-cell morphological metrics to assess the quality of several independent segmentations of a time-lapse confocal movie of a developing starfish embryo, up to the 512-cell stage, with fluorescently labeled cell plasma membranes. Originally, a bio-image analyst segmented the movie using a complex workflow. We demonstrate that running an imported, custom-trained Cellpose 3D model through MorphoNet’s graphical interfaces greatly increases the number of cells that can be successfully segmented.

The third use-case focuses on detecting and resolving regions of under-segmented cells in a confocal 3D dataset of *Arabidopsis thaliana* plant apical shoot meristem, with fluorescent plasma membrane and whole-cell segmentation of 1800 cells. It demonstrates the identification of a large region with heavily under-segmented cells, which is successfully addressed by targeting Cellpose to the specific region of interest.

The fourth use-case addresses the correction of low-quality automated cell nuclei segmentation in a time-lapse confocal *Caenorhabditis elegans* embryo with poor-quality nuclei labeling up to the 350-cell stage, despite accurate manual tracking. It demonstrates how object edition plugins can be used to correct individual nuclei segmentations and propagate these corrections over time.

The final use-case illustrates the efficient detection and correction in the cell lineage of rare residual errors that escaped previous scrutiny in a published time-lapse multiview light-sheet imaging dataset of a developing *Phallusia mammillata* embryo. The dataset features labeled cell membranes and high-quality whole-cell segmentation and tracking. This polishing work is crucial for generating ground truths for studies of natural variation.

All the evaluation and curation steps for the use cases are detailed in the Supp. Mat.

### Use-case 1: Assessing and improving segmentation quality of *Tribolium castaneum* Embryo using unsupervised nuclei intensity metrics and iterative training on partial annotation

This use-case introduces two unsupervised metrics for assessing dataset quality without ground truth, demonstrating their application in evaluating segmentation performance. It also highlights the effectiveness of an iterative segmentation and training approach, where applying a Cellpose model trained on a curated subset significantly improves overall segmentation quality.

#### Dataset description

This dataset features a 59-time-step light-sheet microscopy acquisition of labeled cell nuclei from a developing *Tribolium castaneum* embryo, containing over 7000 cells per time step. It includes two types of ground truth: a *gold truth* (GT), which is expert manual tracking of 181 nuclei over all time steps, and a *silver truth* (ST), generated through a complex pipeline combining the outputs of up to 16 top-performing algorithms selected from 39 submissions. The ST provides automated segmentation masks using a modified label fusion approach for improved accuracy in dense datasets. The GT and ST are independent and do not share common elements.

#### Type of errors

This dataset serves as a benchmark in the fifth Cell Tracking Challenge^6^ to evaluate and compare AI-based image segmentation methods and is therefore regarded as a reliable ground truth. The manual tracking GT is of high quality (expert annotation), but the automated ST is of lesser quality. To evaluate the segmentation quality of the dataset at first time point, the intensity image and the two ground truths (ST and GT) were simultaneously displayed in separate MorphoNet channels (Fig. 3b). Systematic visual inspection identified 12/181 GT points without corresponding nucleus in the ST data; 8 misplaced segmentations where the GT point is not inside the segmentation; 56 over segmentations and many suspicious nuclear shapes.

**Figure 3:**
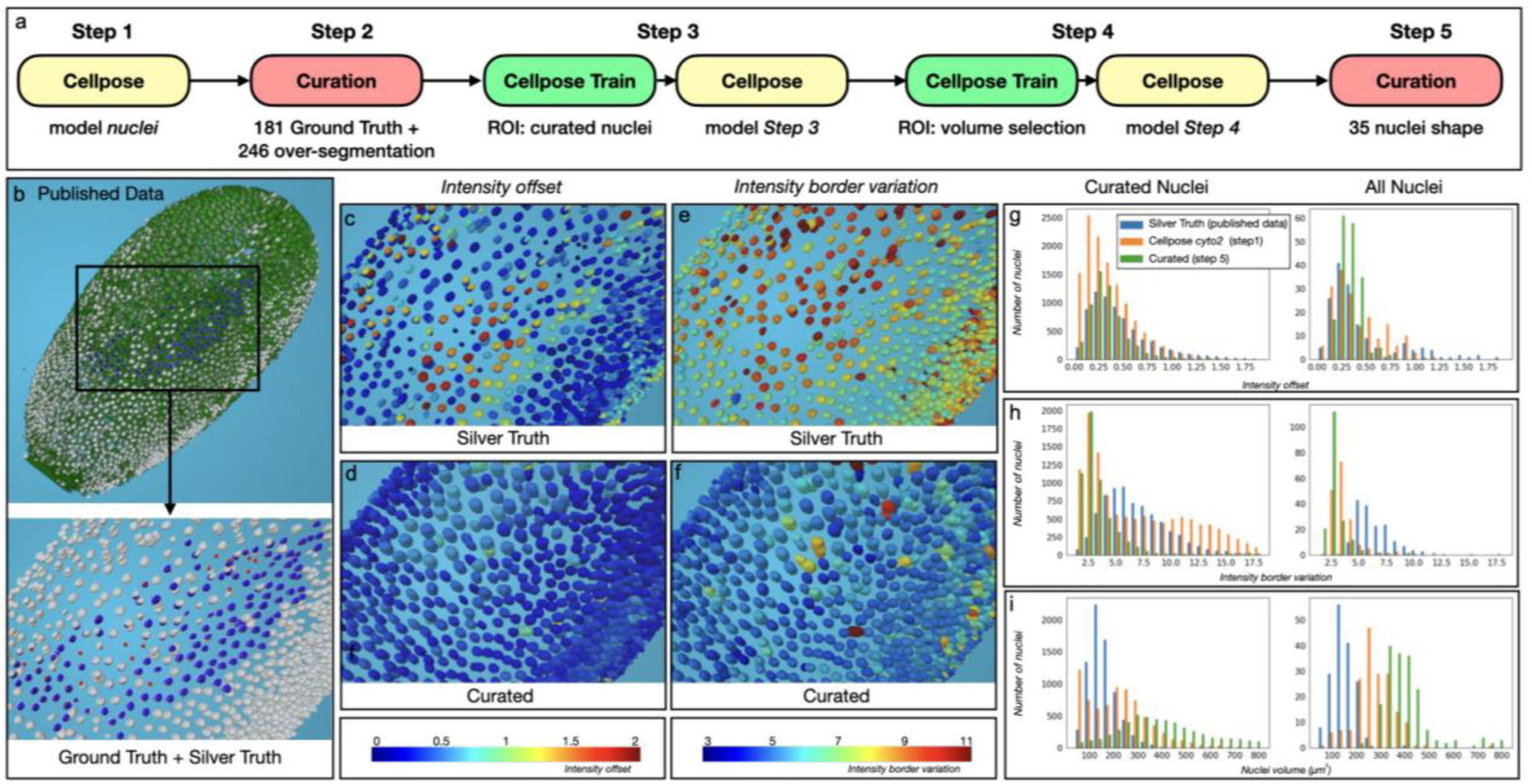
Unsupervised quality assessment and curation of the *Tribolium castaneum* embryo. a. The five steps of the curation pipeline. b. View of the original intensity images (in green) of the first time step of the published data^6^. Both Ground Truth channel (GT, red point) and Silver Truth (ST, white nucleus segmentation) are shown at the top. Blue segmentation corresponds to the match between ST and GT. Bottom, zoom at the GT region (ROI) without the intensity images. c. Projection for the ST of the distance between the gravity center of the intensity inside the segmentation and the centroid of the segmentation. Color bar at the bottom of d. d. Same as c. for the curated pipeline. e. Projection for the ST of the deviation of the intensity at the border of the segmentation. Color bar at the bottom of f. f. Same as e. for the curated pipeline. g. Comparative histogram of the *intensity_offset* property distribution between the ST, the Step 1 and the Step 5 for the 181 curated nuclei (left) and the whole image (right). h. Same as g for the distribution of the *intensity_border_variation* property. i. Same as g for the distribution of the *nuclei volume* property.

#### Error identification

Using the *Match* plugin, each GT position was automatically associated with a corresponding nucleus in the ST, leaving 11 out of 181 positions unassigned (Fig. 3b), indicating approximately 6% of missed nuclei in the ST. To identify inaccurately shaped segmentations, we used two signal intensity-based MorphoNet metrics. The first metric measures the distance between the geometric center of a segmented object and the center of mass of the signal intensity (Supp. Fig. 2 and Supp. Mat. for the full properties description), with 32 of the 56 over-segmented nuclei falling into the upper quartile of this metric’s distribution (Fig 3c-d and Supp. Fig. 3). The second metric evaluates deviations in the signal intensity along the segmentation boundary, with 29 out of 56 over-segmented nuclei identified in the upper quartile (Fig. 3e-f and Supp. Fig. 2). These errors were visualized in 3D by projecting both metrics onto each nucleus, enabling spatial identification of potential segmentation issues (Supp. Fig. 3).

#### Error correction

Due to the high number of errors in the published ST, a de novo nucleus segmentation pipeline was created in MorphoNet with minimal manual intervention. The five-step process (as detailed in the Supp. Mat. UC1 Curated pipeline) included initial segmentation using the standard Cellpose nuclei model, manual curation of 181 GT nuclei, iterative training of a custom Cellpose model, and refinement of segmentation using geometric properties. Key steps involved correcting segmentation errors with various plugins, extending the Cellpose nuclei model using curated data, and refining non-convex shapes. The pipeline significantly improved segmentation quality compared to the published ST and standard Cellpose model (Fig. 3i). Analysis of the curated nuclei revealed a more heterogeneous volume distribution, fewer over-segmentations, and better alignment of intensity-based metrics (Fig. 3g-h), showing substantial improvement over the original segmentation pipeline. We also created a new set of manually segmented ground truth cells (see Supp. Mat. UC1 ground truth protocol), distributed across the ranges of both *intensity_offset* and *intensity_border_variation*. Using this reference, we evaluated the published and curated segmentations (Supp. Fig. 3h–j), confirming that our pipeline both reduces segmentation errors and better matches expert annotations. Moreover, the analysis revealed a clear correlation between the unsupervised metrics and segmentation accuracy. These results validate the utility of our metrics and demonstrate that the iterative pipeline yields higher-quality segmentations than the published dataset.

### Use case 2: Evaluating membrane segmentation quality using *smoothness* metrics and simplifying a segmentation workflow for *Patiria miniata* starfish embryo

This use case thus introduces new unsupervised morphological metrics (the *smoothness* property) to assess segmentation quality and demonstrates that using a custom-trained 3D Cellpose model via graphical interfaces allows extending segmentation to areas of the dataset with lower intensity/noise ratio. Visualization of metrics such as smoothness and cell volumes simplifies the identification of cells that need further manual curation.

#### Dataset description

This dataset comprises 300 3D stacks from time-lapse, two-channel confocal imaging of twelve *Patiria miniata* wild-type and compressed embryos, captured between the 128- and 512-cell stages^23^. Fluorescent labeling highlights cell membranes and nuclei, though imaging was limited to about half of each embryo due to poor penetration and the use of low magnification air objectives. The published whole-cell segmentation (see Supp. Mat. UC2 Original workflow) relied on a complex, multi-step workflow adapted from the CartoCel^24^ requiring advanced bioanalysis expertise. For this use case, a particularly challenging dataset - a compressed wild-type embryo at the 512- cell stage - was selected for evaluation and curation.

#### MorphoNet segmentation with the advanced Cellpose plugin

We explored Cellpose’s potential to improve segmentation quality in challenging regions of a 3D dataset. Initial segmentation using the Cyto2 model identified 1187 cells, with a bimodal size distribution and 939 cells smaller than 1000 voxels (Fig. 4i). The Deli plugin, which eliminates these small cells (Fig. 4j), brought the number of cells down to 211 (Fig. 4j); however, analysis with the *smoothness* property (Fig. 4g) revealed rough cell surfaces. To address this, we further extend the training of the *Cyto2* model on the published 3D database^23^ (see Supp. Mat. UC2 Training Protocol workflow). The retrained model produced 284 cells with again a bimodal size distribution (Fig. 4i) which was corrected with the *Deli* plugin. This significantly simpler pipeline produced smooth cell segmentations (Fig. 4h), with a size distribution comparable to the published ground truth and a high Intersection over Union (IoU) score (Supp. Fig. 5). While two cells were missing, and 11 under-segmented compared to the published data (Fig. 4f), this new approach recovered 48 additional cells (Fig. 4e). while simplifying the segmentation process compared to the original pipeline.

**Figure 4:**
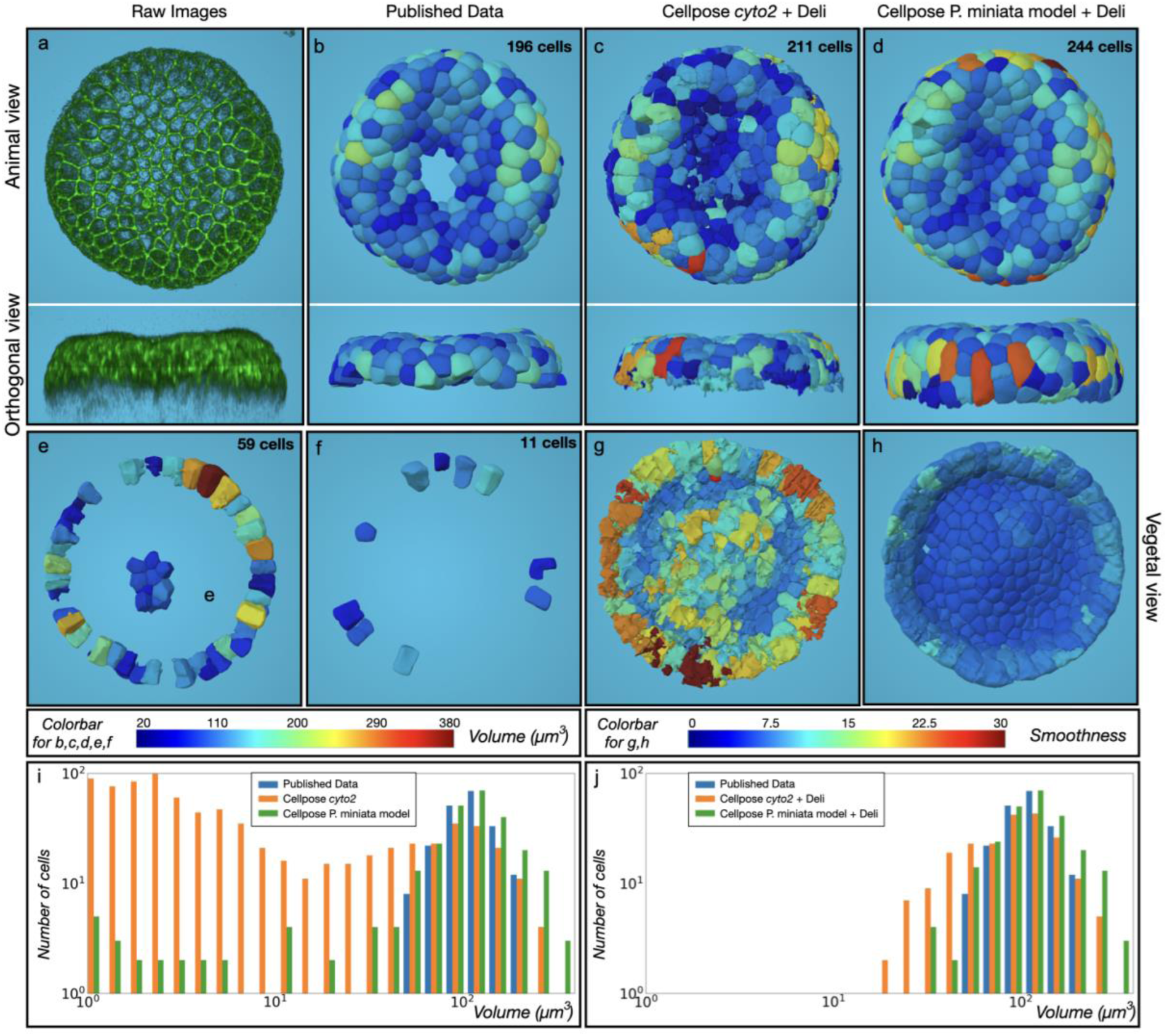
Starfish whole-cell segmentation using Cellpose. *a. Animal and Lateral view of the maximum intensity projection from the published dataset^23^*. *b. Segmented ground truth image*. *c. Result of the segmentation using Cellpose Cyto2 model followed by the removal of small cells (<1000 voxels) with the Deli plugin* *d. Result of a Cellpose segmentation with a model trained on P. miniata dataset followed by the Deli plugin*. *e. New cells created (different between b and d)* *f. Under segmentation and missing cells generated by d. compared to b*. *g. Vegetal view of c with smoothness representation*. *h. Opposite view of d with smoothness representation*. *i. Cell size distribution in the published segmentation (b), Cellpose cyto2 model (c) and Cellpose with P. miniata model (d)*. *j. Identical as i after application of the Deli plugin*. *Colors in b-f, represent cell volume in µm^3^*. *Colors in g,h represent Smoothness*.

### Use case 3: Interactive targeted segmentation using CellPose for resolving under-segmentation in *Arabidopsis thaliana* shoot apical meristem whole-cell segmentation

This use case demonstrates the power of interactive selection with targeted segmentation methods that can resolve under-segmentation issues in complex datasets.

#### Dataset description

This dataset^25^ consists of a 19-time steps time-lapse confocal acquisition of a live *Arabidopsis thaliana* shoot apical meristem with fluorescently-labeled cell membranes. Cell numbers ranged from 600 cells at the first time point to around 1800 cells at the last (Fig. 5a). We use the last time step (19) with the higher number of cells to illustrate the curation procedure.

**Figure 5:**
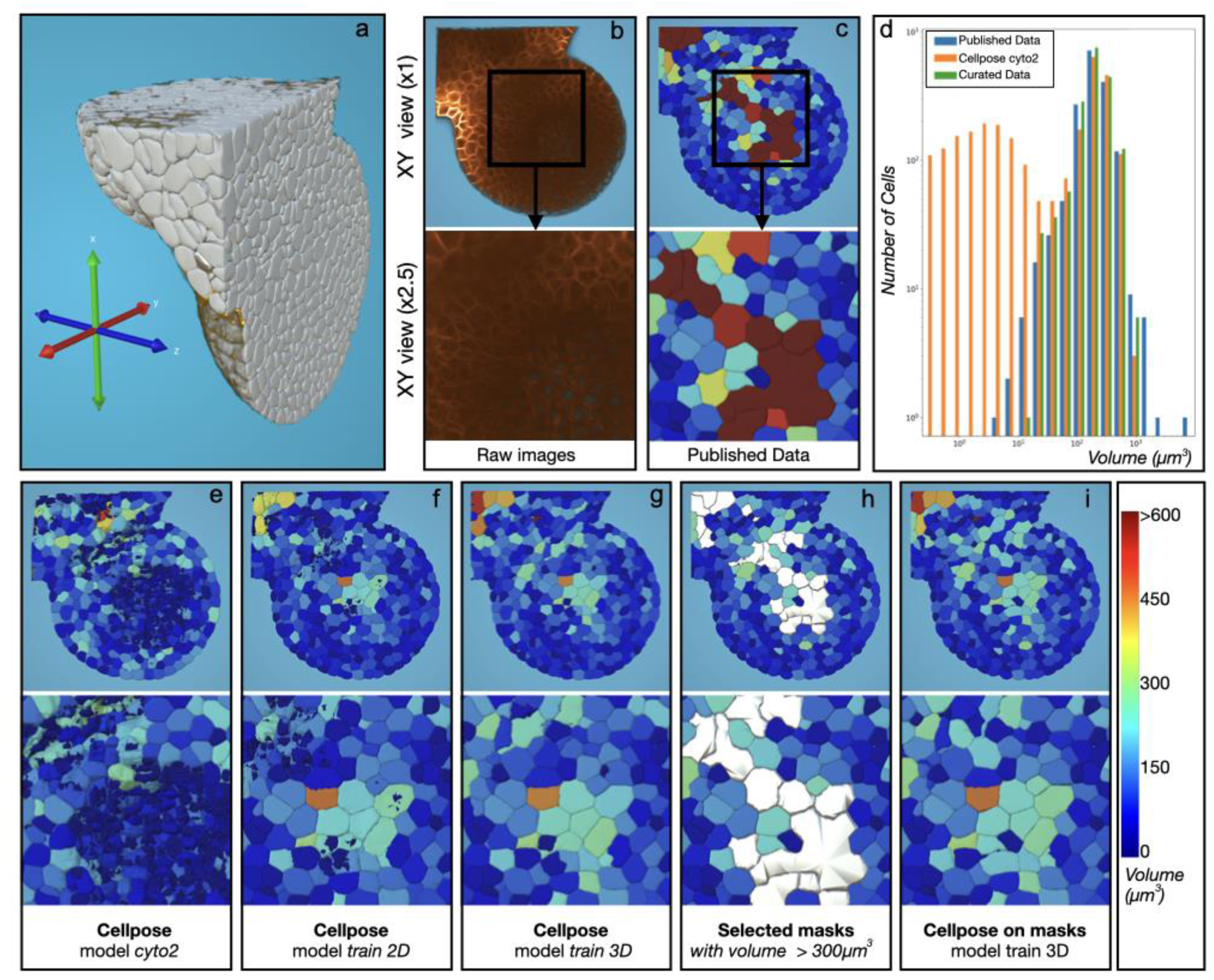
Curation of a segmented shoot apical meristem of *Arabidopsis thaliana*. *Visualization of the time step n°19 for several types of curations using MorphoNet*. *a. 3D view of an Arabidopsis thaliana shoot apical meristem^25^*. *b. Published 3D intensity images*. *c. Published 3D segmentation of an Arabidopsis thaliana shoot apical meristem^25^ obtained using MARS-ALT^26^*. *d. Comparative histogram based on the cell volume between the published segmentation (c), the result of the cyto2 prediction (e) and the final curated version* *(i) + Deli plugin for cells <1000 voxels. X and Y axes are in log scale*. *e. Result of the Cellpose^29^ 3D MorphoNet plugin using the pretrained cyto2 model*. *f. Result using the Cellpose 3D MorphoNet plugin using the model trained over the 10 first time steps with the Cellpose training plugin with the XY planes*. *g. Result of the Cellpose 3D MorphoNet plugin using the model trained over the 10 first time steps with the Cellpose training plugin with each plane of the 3D images*. *h. Selected masks larger than 300µm^3^ in the published dataset (c)*. *i. Result of the Cellpose 3D MorphoNet on the selected masks (h) using the model trained over the ten first time steps with the Cellpose training plugin with each plane of the 3D images*.

#### Error detection

The analysis of the published segmentation uncovered multiple under- and over-segmentation errors in the deep cell layers, caused by poor image quality in the inner regions of the meristem (Fig. 5b). These errors hindered accurate cell tracking over time. Observing the size homogeneity of shoot apical meristem cells (Fig. 5d), we hypothesized that larger-than-expected cells might indicate under-segmentation errors, while smaller cells could result from over-segmentation. By projecting the automatically computed cell volumes as a color map onto the segmentations, we identified a large under-segmented region containing 20 cells in the deep meristem layer (highlighted in red in Fig. 5c), along with a few small over-segmented cells (Fig. 5i).

#### Error correction

The large number of fused cells in the under-segmented region prompted us to test whether our interactive Cellpose plugin could outperform the original MARS/ALT software^26^. The standard pretrained *cyto2* model of Cellpose^2^ produced numerous very small over-segmented cells (Fig. 5d and Supp. Fig. 6), suggesting it was not well-suited for this dataset^27^. To improve performance, we used the high-quality MARS/ALT segmentations from the first ten time points and employed our Cellpose train plugin to extend the *cyto2* model training. We tested two training modalities: “2D” training, using only the XY planes, and “3D” training, incorporating XY, XZ, and YZ planes. The segmentations after additional 2D training still contained many over-segmented cells (Fig. 5f), a problem that was significantly reduced with 3D training (Fig. 5g). Despite this, the heterogeneity of signal intensities still caused over-segmentation in more superficial regions of the meristem. We thus targeted the Cellpose plugin only to the large under-segmented region (Fig. 5h) and removed small over-segmented cells using the *Deli* segmentation correction plugin (Fig. 5i). This targeted approach was fast and accurate (Fig. 5d) and allowed us to preserve the accurate segmentations from the original published data while curating the identified errors. Notably, the curated result restored a unimodal and symmetric volume distribution (Supp. Fig. 6g), consistent with the expected homogeneity of the shoot apical meristem^28^, which was further confirmed by visual inspection (Supp. Fig. 6h-i). With this approach, we successfully curated 20 under-segmented regions and generated 98 new cells.

### Use-case 4: Improving segmentation quality using object editing plugins for Caenorhabditis elegans cell lineage

This use case illustrates how object editing plugins can be employed to improve segmentation quality in challenging datasets despite poor-quality nuclei labeling.

#### Dataset description

This dataset^30^ consists of a live 3D time lapse confocal acquisition of a developing *Caenorhabditis elegans* embryo with fluorescently-labeled cell nuclei (Fig. 6a), over 195 time steps between the 4- and 362-cell stages. This dataset was included in the Cell Tracking Challenge^6^ (see the Supp. Mat. UC4 for the original workflow).

**Figure 6:**
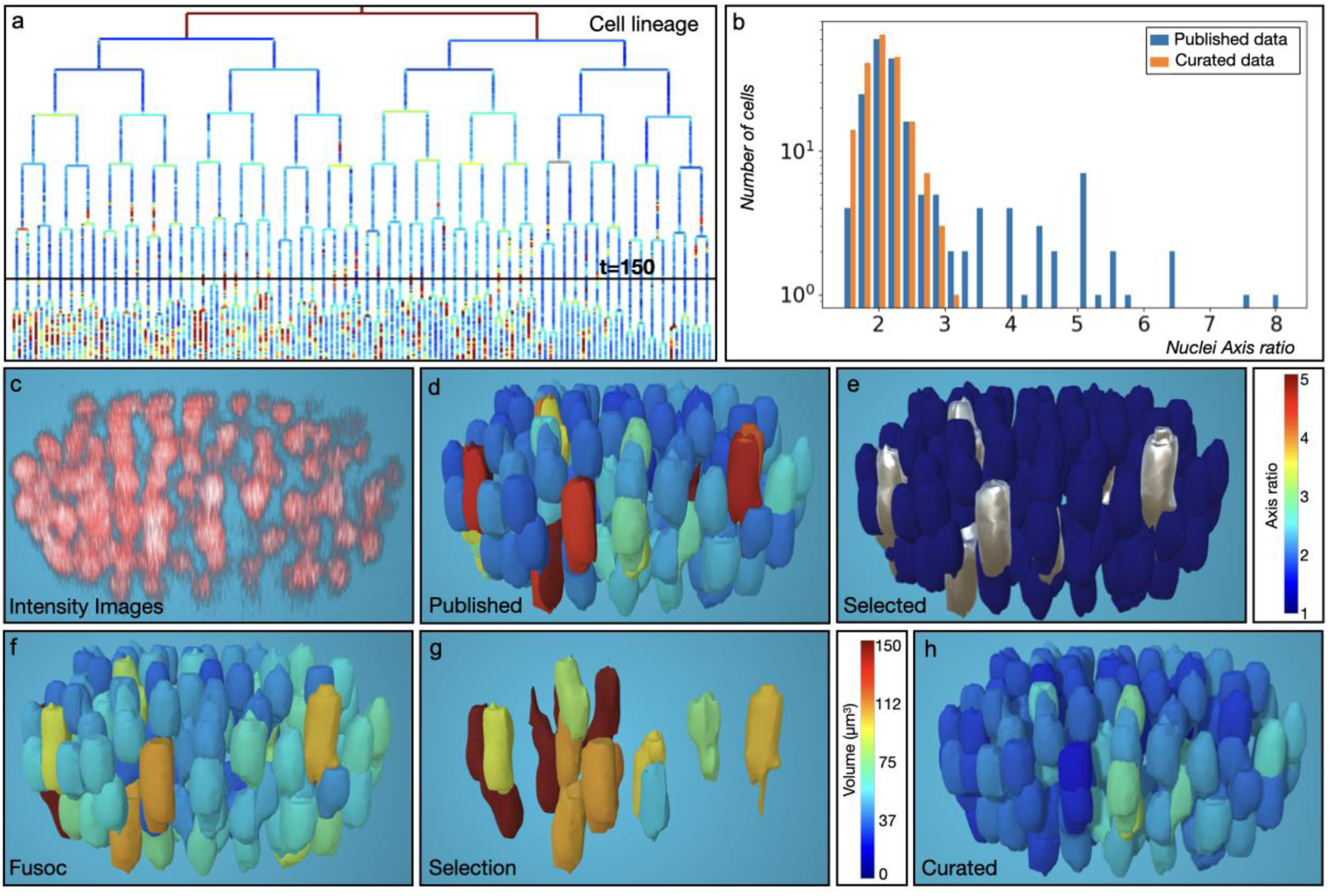
Curation of a *Caenorhabditis elegans* embryo dataset. *a. Cell lineage viewer of all time steps colored by clonal cells of the published segmented dataset^30^*. *b. Comparative histogram based on the axis ratio between the published data and the curation*. *c. 3D View of the intensity images at t=150*. *d. 3D View of the published segmented dataset at t=150. Colors represent the ratio of the longest axis on the shortest axis of the shape*. *e. Automatic selection (in gray) of the nuclei with an axis ratio>2.9*. *f. Result of the Fusoc plugin applied on all selected nuclei. Colors represent the nuclei volume in µm^3^*. *g. Same as f. But only previously selected nuclei are shown*. *h. Result of the Gaumi plugin applied independently on regions fused with four, three or two nuclei. Colors represent the axis ratio as in d*.

#### Error detection

Although the manually curated published cell lineage appeared accurate (Fig. 6a), some touching nuclei were elongated with flat vertical contact interfaces, likely due to poor imaging quality. To identify these segmentation artifacts systematically, we calculated the axis ratio for each segmented object and visualized these values directly on their surface meshes (Fig. 6b). The distribution revealed two peaks: a sharp one around 2, containing most nuclei, and a broader one above 3. Nuclei with an axis ratio greater than 2.9 were flagged for potential correction.

#### Error correction

We curated these errors for a specific time point by first applying a fusion correction plugin (Fig. 6f and g) to all selected nuclei pairs with an axis ratio greater than 2.9 (Fig. 6e). This was followed by a separation plugin (Fig. 6h) that divides fused objects into distinct nuclei using a Gaussian mixture applied to intensity values. We initially addressed the more complex cases manually, which included one group of six fused nuclei, one group of four fused nuclei, and two groups of three. Next, we automatically curated the remaining nine simpler cases of two fused nuclei by selecting nuclei with a volume greater than 80 µm³. As expected, the axis ratio distribution of the curated set became unimodal and centered around 2 (Fig. 6b), reflecting more regular nuclear shapes—an improvement confirmed by visual inspection (Supp. Fig. 7). The entire time step at t=150 was fully curated with just 5 plugin actions (Fig. 6h).

### Use-case 5: Enhancing cell lineage accuracy by detecting segmentation errors in *Phallusia mammillata* embryos

This use case showcases how MorphoNet can efficiently detect and correct segmentation errors in complex 3D+t datasets of developing ascidian embryos, enhancing the accuracy of cell lineage reconstructions and analysis of cell division timing variations.

#### Dataset description

This dataset^16^ comprises 64 time steps of multi-view light-sheet microscopy of developing ascidian embryos (64–298-cell stages), segmented using the ASTEC algorithm^16^. Despite the high-quality segmentation and tracking, residual errors such as multi-component cells, over- and under-segmentations, delayed or missed divisions, and shape inaccuracies remained, with no tools available at the time of the publication to systematically validate or correct the 10,000 cell snapshots.

#### Error detection

MorphoNet was used to polish the cell lineage to a sufficient level to study natural variation in cell division timing. The main challenge was identifying rare errors within a cell lineage that appeared accurate by visual inspection. To address common lineage residual errors such as missing or delayed divisions and broken lineage links, we developed a set of segmentation error identification metrics. These metrics were visualized by projecting their results onto the lineage using an interactive viewer linked to the 3D dataset (Fig. 7a and b). All identified errors stemmed from segmentation issues rather than tracking, and their correction was streamlined by the bidirectional connection between the lineage and embryo representation windows, allowing direct navigation between the two.

**Figure 7:**
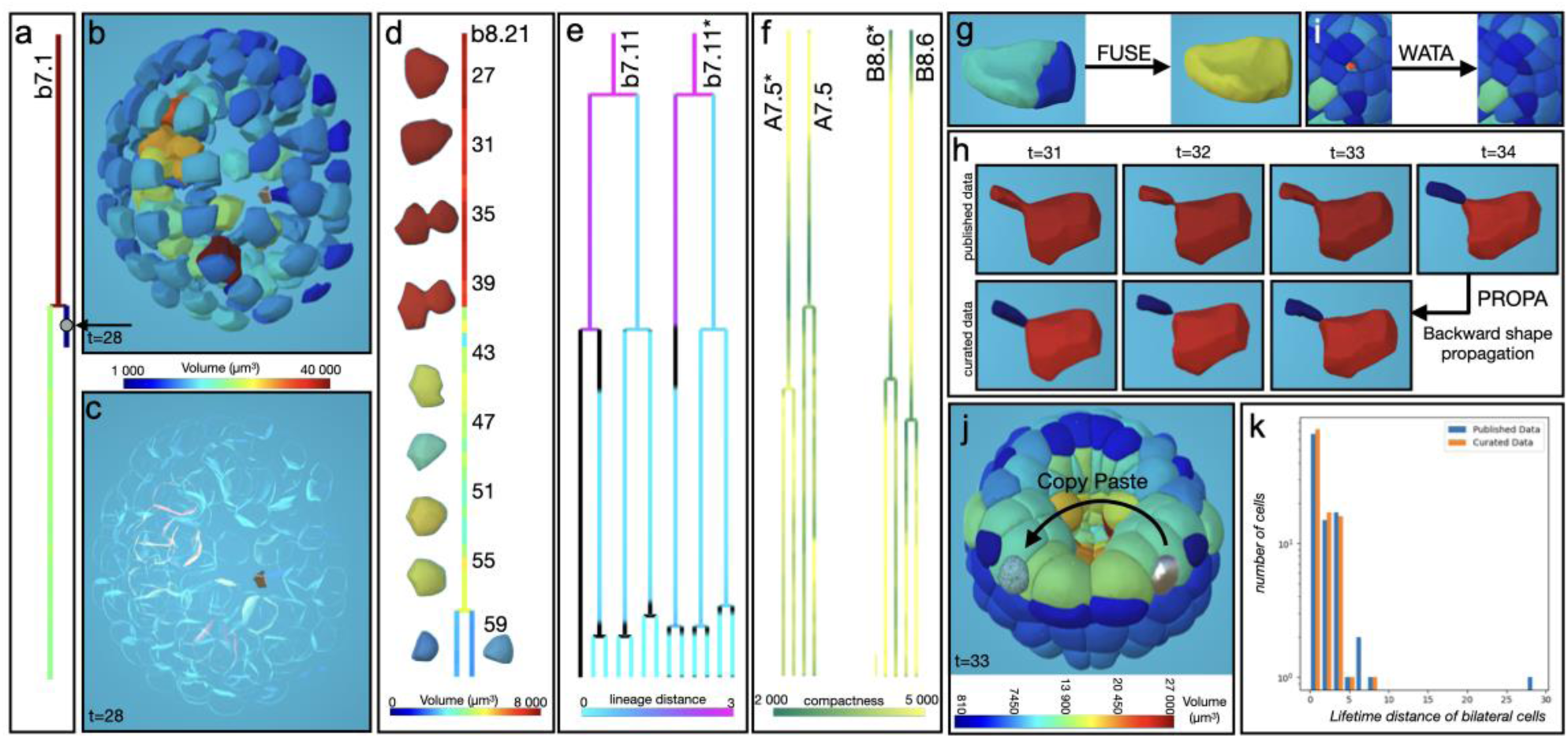
Curation of a Phallusia mammillata ascidian embryo named Astec-Pm9 in the publication^16^. a. Cell lineage of the cell b7.1 after the execution of the Disco and Deli plugins on the published data. Projection of the volume property, the *colormap between b and c represents the cell volume in µm^3^*. b. *Scatter view of the corresponding segmented embryo described in a. at t=28 colored by cell volume. Colormap identical as a*. c. *Same view as b. with the activation of the “highlight” mode which focuses on the selected cell and shows other cells in transparent colors*. d. *Several snapshots of the cell b8.21 at different time points with its associated cell lineage. Colormap represents the cell volume in µm^3^ which points to a missing division*. e. *Cell lineage of the bilateral cells b7.11 and b7.11*. Colorbar shows the lineage distance between the bilateral symmetrical cells. Black region represents snapshots with no matches between bilateral symmetrical cells*. f. *Cell lineage of the bilateral cells a7.5 and b8.6. Colorbar shows the compactness property. The property highlights that the delay of division between A7.5* and A7.5 is due a under-segmented error of A7.5*. B8.6 and B8.6* have expected behavior*. g. *Result of the Fuse plugin applied on an over-segmented cell*. h. *Top line: Several snapshots of B7.7* cell under-segmented (between time point 31 and 33) from the original segmented embryo. Bottom: result of the Propa plugin applied backward from time t=34 where both cells are well separated*. i. *Example of the result of the Wata plugin from a manual added seed (red dot) in the empty space*. j. *Result of the Copy-Paste plugin from the selected cell (in gray) on the left side of the embryo to the right side (the new cell appears with a mesh shader in blue). Colormap represents cell volume in µm^3^*. k. *Comparison of the lifetime of bilateral symmetrical cells. The X axis shows the number of in time points separating the division of bilateral cell pairs. The Y axis corresponds to the number of cell (in log)*

State-of-the-art segmentation methods often produce systematic errors, such as very small segmented objects, which we identified using geometric properties like cell volume (e.g., cell a7.1, Fig. 7a). Property projection, combined with scatter visualization, facilitates the identification of cells even in dense, interior regions (Fig. 7b), while the highlight mode enhances this process by isolating the cell of interest (Fig. 7c). Occasional missing or false divisions disrupt accurate cell histories. During early ascidian embryogenesis, stable cell volumes allowed volume projections onto the lineage to identify rapid variations, revealing missed divisions (Fig. 7d).

Ascidian embryos exhibit bilateral symmetry, with homologous cells dividing almost synchronously. A l*ineage distance* property^16^ revealed missed divisions (e.g., b7.11* in Fig. 7e). To distinguish segmentation errors from natural variability, a *compactness* property was used to detect inaccuracies in division timing by tracking cell rounding during mitosis (e.g., Fig. 7f).

Using MorphoNet, an expert identified approximately 20 isolated cell lineage errors in a dataset of 185 cell snapshots in just two hours.

#### Error correction

The 41 multi-component labels were separated using the *Disco* plugin, identifying seven potential cells via the *volume* property analysis (Fig. 7b). The others, considered as small artifacts, were merged with neighboring cells sharing the largest surface area using the *Deli* plugin. Division timing issues often stem from over-segmentation, under-segmentation, or missing cells. Over-segmentations were resolved with the *Fuse* plugin (Fig. 7g), while under-segmentations were corrected using *Propagate segmentation* plugins by tracing back from the first accurate segmentation of sister cells to their mother’s division point (Fig. 7h). For missing cells, seeds were added with seed generator plugins and segmented using local seeded watershed algorithms (Fig. 7i). If imaging quality was too poor, the bilaterally symmetrical cell served as a mirror-image proxy, using the Copy-Paste plugin to replicate, rotate, and scale the cell to the missing position (Fig. 7j). A total of 185 errors were corrected in approximately eight hours using 264 MorphoNet plugin actions, resulting in a more accurate estimation of natural variability in cell division timing (Fig. 7k and Supp. Fig. 8). This duration includes the full workflow performed by a single user: loading the dataset, exploring the cell lineage, identifying segmentation or division errors, and applying corrective plugins. The reported actions include both exploratory attempts and validated corrections, combining user interaction with computation time.

## Discussion

Recent advancements in optical time-lapse microscopy allow for the 3D capture of dynamic biological processes but face challenges in automating the segmentation and tracking of large, complex datasets. Residual segmentation errors in time-lapse datasets disrupt data interpretation and hinder long-term cell tracking. Moreover, accurate, high-quality 3D data is critical for training next-generation AI-based segmentation tools, yet the availability of such datasets remains limited.

To address these challenges, **MorphoNet 2.0** was developed as a standalone application for reconstructing, evaluating, and curating 3D and 3D+t datasets. Building on its previous web-based version, MorphoNet 2.0 integrates voxel images with meshed representations, leveraging both Unity Game Engine and Python for enhanced interactivity and processing power. Key features include tools for assessing segmentation quality through unsupervised metrics (e.g., volume, smoothness, and temporal stability), automated error detection, and visualization of segmented objects alongside raw intensity images.

Conventional image curation tools struggle with the complexities of interacting with 3D voxel images. In contrast, MorphoNet 2.0 introduces a novel approach using dual representation layers, enabling efficient biocuration by isolating problem areas and significantly accelerating dataset refinement. MorphoNet 2.0 is user-friendly, open-source, and supports both scientific discovery and AI training by producing high-precision 3D datasets and enabling reproducible, scalable data analysis.

By showcasing five use-cases of fluorescence datasets in which cell membranes or nuclei were labeled, we demonstrated that the tool’s high versatility and user-friendliness enables biologists without programming skills to efficiently and intuitively detect and handle a broad range of errors (under-segmentation, over-segmentation, missing objects, lineage errors).

MorphoNet has a high potential to adapt to evolving datasets and segmentation challenges. First, its open-source Python plugin architecture fosters community-driven improvements. These could target cellular datasets as exemplified by the five use-cases presented. They could also open MorphoNet to other imaging modalities, including multi-modal datasets combining for instance fluorescence and electron microscopy. Additional plugins could accommodate new AI tools or automate the training of segmentation models, through data augmentation, feature selection, or hyperparameter optimization. Plugins for widely used platforms such as Napari or Fiji will also broaden the user-base and interoperability of the tool, as would also the development of flexible export options to integrate MorphoNet outputs with other analytical pipelines or visualization software. The MorphoNet platform could be further extended through the creation of a centralized repository for community-developed plugins, the organization of MorphoNet-based bio-image analysis challenges to stimulate community engagement, and the provision of curated datasets to serve as benchmarks for testing and validating new segmentation algorithms. To support these developments over time, we rely on institutional support from our host laboratories and research organizations, and we will seek additional funding through dedicated research grants. We also aim to foster open-source contributions, develop training materials, and provide user support to ensure long-term adoption and sustainability.

Curation could also be improved by the introduction of cloud-based, multi-user capabilities to enable experts to work on the same dataset simultaneously. For now, web browsers impose constraints on the use of computing resources, which could be lifted in the near future, for example through the development of WebGPU.

Curation capabilities will likely also be enhanced by the implementation of adaptive machine learning models that integrate past user corrections to suggest or automate future edits. In the context of 3D segmentation tasks, manual expertise and curation for voxel-wise segmentation is labor-intensive and expensive. Full annotation of large datasets may thus not be feasible. Partial annotations allow datasets to be created with less time and resources while still providing valuable information. Including algorithms such as Sketchpose^31^ could leverage weakly-supervised learning to generalize from the partially annotated data and infer segmentation patterns in the unlabeled portions of the dataset, a strategy we initiated with the Tribolium dataset. This will make it possible to train models on diverse datasets without the burden of full annotations.

By addressing major challenges in 3D and 3D+t dataset assessment and curation, MorphoNet 2.0 provides a versatile platform for improving segmentation quality and generating reliable ground truths. Its user-friendliness, adaptability and extensibility position it as a valuable tool for advancing quantitative bio-image analysis, with significant potential for enhancement and broader application in the future.

## Acknowledgements

This work was supported by grants of the Occitanie Region (ESR-PREMAT-213) and of the French National Infrastructure France BioImaging (ANR-10-INBS-04) to EF, and by the Cell-Whisper (ANR-19-CE13-0020) and scEmbryo-Mech (ANR-21-CE13-0046) ANR projects coordinated by PL. EF and PL were CNRS staff scientists. NF was an assistant professor at UM. KB was a UM Phd Student supported by the EpiGenMed Labex (ProjetIA-10-LABX-0012) and a post-doc funded by the scEmbryo-Mech project. BG,TL were contract CNRS engineers. AC was a UM Master student. We thank Christophe Godin, Vanessa Barone,Thibault de Villèle and Volker Baecker for their valuable feedback and advice.

## Author Contributions

B.G. contributed to the development of the website, the 3D viewer and the python API. TL contributed to the development of the 3D viewer and the python API. KB contributed to its featuring and to the help pages. AC and NF contributed to the *Deform* plugin. P.L. contributed to the development of features and to writing the manuscript. E.F. conceived the concept of MorphoNet, developed the code, contributed to its featuring and wrote the manuscript. All authors contributed to discussions on the structure of the manuscript.

