## Supplementary Materials for "MorphoNet 2.0: An innovative approach for qualitative assessment and segmentation curation of large-scale 3D time-lapse imaging datasets"

| Datasets | Dataset 1 : Caenorhabditis elegans (timelapse of nuclei channel) | Dataset 2 : Phallusia mammillata (membrane + nuclei channels) | Dataset 3 : Phallusia mammillata (timelapse of membrane channel) | Dataset 4 : Tribolium confusum (timelaps of nuclei channel) |
|---|---|---|---|---|
| image types | Segmented and intensity images | Intensity image only | Segmented and intensity images | Segmented and intensity images |
| image dimensions (x,y,z) | 708*512*35 | 737*550*518 | 732*988*1084 | 965*1871*991 |
| time points | 40 time points | 1 time point | 21 time points | 10 time points |
| number of cells | from 8 to 26 per time point | None (no segmentation) | from 64 to 215 per time point | around 7000 per time point |

| Computer OS | Computer type | Memory (RAM) | Processor (CPU) | Graphic Card (GPU) | Dataset 1 | Dataset 2 | Dataset 3 | Dataset 4 | Diagnosis |
|---|---|---|---|---|---|---|---|---|---|
| MacOS Silicon (Sonoma 14.6.1) | Laptop (60 fps cap) | 64 Gb | M1 Max | M1 Max | 111 fps Great | 100-110 fps Great | 95-111 fps Great | 42-60 fps Great, but slow | The machine is perfectly adapted to the task |
| MacOs Big Sur (11.7.10) | Laptop (60 fps cap) | 16 Gb | i7 4 cores 2,2 GHz | Intel iris pro (No dedicated graphics card) | 5 fps Almost impossible | 3 fps Impossible | 3 fps Impossible | 3 fps Impossible | The machine is only suited for datasets without intensity images, and is ill suited for curation. It would benefit from a graphics card |
| Windows 11 | Laptop (144 fps cap) | 16 Gb | i7 8750H 2,2GHz 6 Cores 12 threads | NVIDIA GeForce RTX 2070 Laptop | 144 fps Great | 100-144 fps Great | 120-144 fps Great | 3-30 fps Very difficult | The machine is fine for most cases, but will struggle on large datasets, held back by the amount of RAM |
| MacOS Silicon (Ventura 13.3.1) | Desktop (60 fps cap) | 16 Gb | Apple M2 | Apple M2 | 60 fps Great | 30-60 fps Good | 30-40 fps Good | 25-55 fps Acceptable, but slow | The machine is fine for most cases, but will be slower on large datasets, held back by the amount of RAM |
| Windows 10 | Laptop (60 fps cap) | 64 Gb | i7-11850H 2,50GHz 8 cores 16 threads | NVIDIA GeForce RTX 3080 laptop | 60 fps Great | 60 fps Great | 60 fps Great | 60-30 fps Great, but slow | The machine is perfectly adapted to the task |
| Ubuntu 20.04.2 LTS | Desktop (60 fps cap) | 64 Gb | Xeon CPU E5-2620 v2 2.10GHz | NVIDIA GV100 TITAN V | 60 fps Great | 60 fps Great | 60 fps Great | 30-60 fps acceptable | The machine is perfectly adapted to the task |
| Ubuntu 20.04 LTS | Desktop (60 fps cap) | 188 Gb | Xeon 20 cores 40 threads 2.1 GHz | NVIDIA Quadro RTX 4000 | 60 fps Great | 60 fps Great | 60 fps Great | 20-50 fps Acceptable | The machine is perfectly adapted to the task. |
| Ubuntu 22.40 LTS | Laptop (60 fps cap) | 64 Gb | Intel i9 2,6 GHz 16 cores | Mesa Intel® UHD Graphics (No dedicated graphics card) | 8 fps Almost impossible | 12 fps Almost impossible | 4 fps Impossible | 4 fps Impossible | The machine is only suited for datasets without intensity images. The specs are great except for the lack of a graphics card. Adding one would make it perfect for the task |
| Ubuntu 24.04.1 LTS | Desktop (120 fps cap) | 64 Gb | Intel Core™ i9-14900K × 32 | NVIDIA GeForce RTX™ 4080 SUPER | 120 fps Great | 120 fps Great | 120 fps Great | 110-120 fps Great | This machine gets remarkable performance all around, The best kind you can get |

**Supplementary Table 1:** *Performance of the MorphoNet standalone application of various devices. The upper table describes the datasets used for benchmarking the application. The lower table gives an evaluation of performances for different devices, as well as a description of the devices specifications. The cells are colored according to whether or not a particular device is adapted for the visualization and/or curation of a particular dataset. The datasets can be accessed at the link below:*

*https://seafile.lirmm.fr/published/morphonet_data/BENCHMARKING_DATASETS*

| I am: | I want to: | Related documentation: |
|---|---|---|
| A biologist | Visualize a dataset **located on the MorphoNet servers** (public or private) | 1. You can either :<br>– Install the standalone app (handles large datasets)<br>OR – Use the website (handles small datasets)<br>2. Use the MorphoNet 3D Viewer |
| | Visualize a dataset with **segmented** and/or **intensity** images on my computer | 1. Create and view local datasets on MorphoNet standalone<br>2. Use the MorphoNet 3D Viewer |
| | Curate a dataset with **segmented** and optionally **intensity** images on my computer | 1. Create and view local datasets on MorphoNet standalone<br>2. Look at a curation example<br>3. Have a look at all available default plugins<br>4. List of automatically computed image properties<br>5. Use the MorphoNet 3D Viewer |
| | Create and segment a dataset with **intensity images** only | 1. Create a local datasets from your intensity images only on MorphoNet standalone<br>2. Use the MorphoNet 3D Viewer |
| | Upload a dataset with **segmented** and/or **intensity** images on the MorphoNet server | You can either :<br>– Upload a dataset from the MorphoNet application<br>OR. – Upload a dataset with the FIJI plugin |
| | Upload a dataset with **meshes** | 1. Look first at the required mesh format<br>2. Upload a dataset with meshes with the FIJI plugin |
| | **Share** a dataset on MorphoNet | Share a dataset on the website |

| I am: | I want to: | Related documentation: |
|---|---|---|
| | Add **properties** to your local dataset | 1. Have a look at the Properties format<br>2. You can use the MorphoNet 3D Viewer to add a property from the menu |
| | Add **properties** to the dataset located on the MorphoNet server | 1. Have a look at the Properties format<br>2. Upload properties with th FIJI plugin |
| | Add and visualize **genetic** properties on a dataset | 1. Create a genetic property<br>2. Add them using the MorphoNet 3D Viewer<br>3. and then use them in Genetic menu |
| | **Download** a full dataset from a dataset | Download the dataset from MorphoNet |
| | Visualize and Interact with a dataset in **Virtual Reality** | 1. Install the standalone app on Windows<br>2. Open the Virtual Reality mode |

| | | |
|---|---|---|
| A **bio-image analyst** | Upload a dataset with **segmented** and/or **intensity** images on the MorphoNet server | Using the API you can convert your segmented images in meshes and then directly upload the meshes . You can look at this example from the documentation |
| | Upload a dataset with **meshes** | 1. Look first at the required mesh format<br>2. Upload the meshes using the API . You can look at this example from the documentation |
| | **Share** a dataset on MorphoNet | Share a dataset with the python API |
| | Add **properties** to the dataset located on the MorphoNet server | 1. Have a look at the Properties format<br>2. Upload properties with the python API |
| | **Download** a full dataset from a dataset | Download the meshes and the properties with the python API |
| | Create custom **plugins** for MorphoNet | 1. Look how to use python API with the Standalone application in developer mode.<br>2. Create custom plugins for bio-curation |
| | Use MorphoNet to **analyze** 3D + t images | Look at the MorphoNet tutorials in python :<br>- Visualize/Analyze complex 3D+time.<br>- How to segment and track 3D cells.<br>- How to create a simple simulation.<br>- How cell adjacency relationships impact cell state transitions. |

| I am: | I want to: | Related documentation: |
|---|---|---|
| A **developer** | Contribute to the development of **MorphoNet 3D viewer** | Clone the MorphoNet Unity GitLab Project |
| | Contribute to the development of **MorphoNet python API** | Clone the MorphoNet Python Project |
| | Integrate MorphoNet into your **website** | Look at the API REST to configure your own urls |

**Supplementary Table 2:** *MorphoNet Documentation hub on various help pages depending on user types and use cases.*

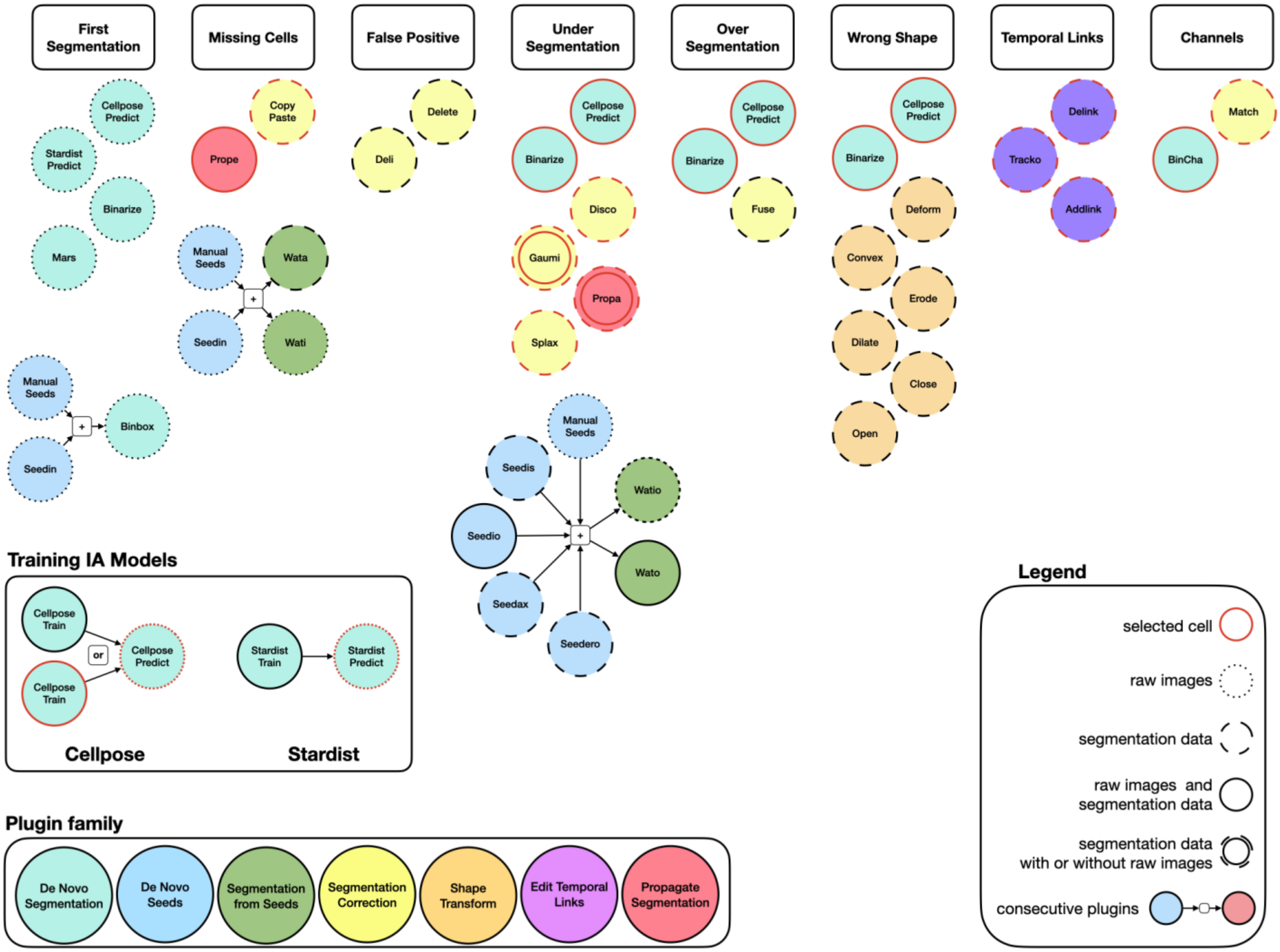


**Supplementary Figure 1: Representation of the MorphoNet Plugins organized by the corresponding functionalities according segmentation issues.**

Color code represents the plugin family. Description of each plugin functionalities is accessible on the help web page : https://morphonet.org/help_curation#plugin_list

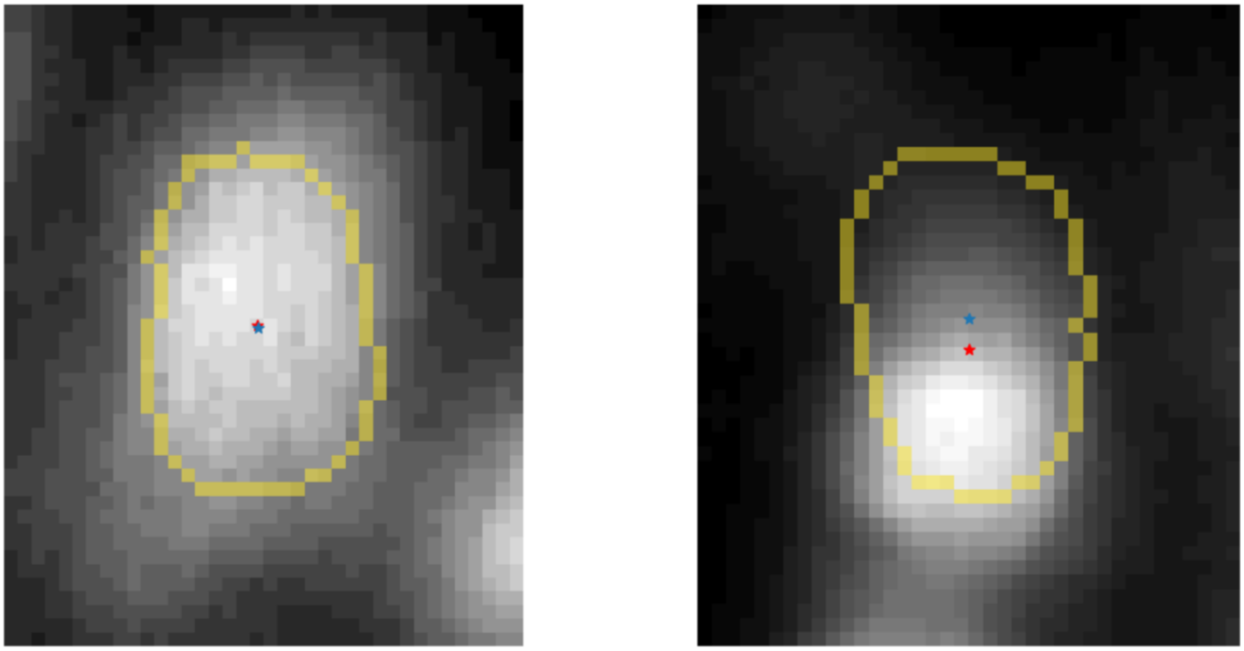

**Supplementary Figure 2: Illustration of properties *intensity_border_variation* and *intensity_offset for 2 nuclei.***

*2D slice of the intensity image of 2 distinct nuclei superimposed with their corresponding segmentation border (in yellow). The blue cross is the corresponding gravity center of the intensity images inside the segmentation. The blue cross corresponds to the geometrical center of the segmentation shape. The intensity_offset is the euclidean distance between the geometric center of the segmented object and the center of mass of the signal intensity. The intensity_border_variation is the standard deviation of the intensity images only at the border of the segmentation (in yellow). The left nucleus, representing a well-segmented example, exhibits low values for both properties (intensity_offset=0.18 and intensity_border_variation=1.84). The right nucleus, where the segmentation is misaligned, shows high values for both properties. (intensity_offset=2.06 and intensity_border_variation=10.73).*

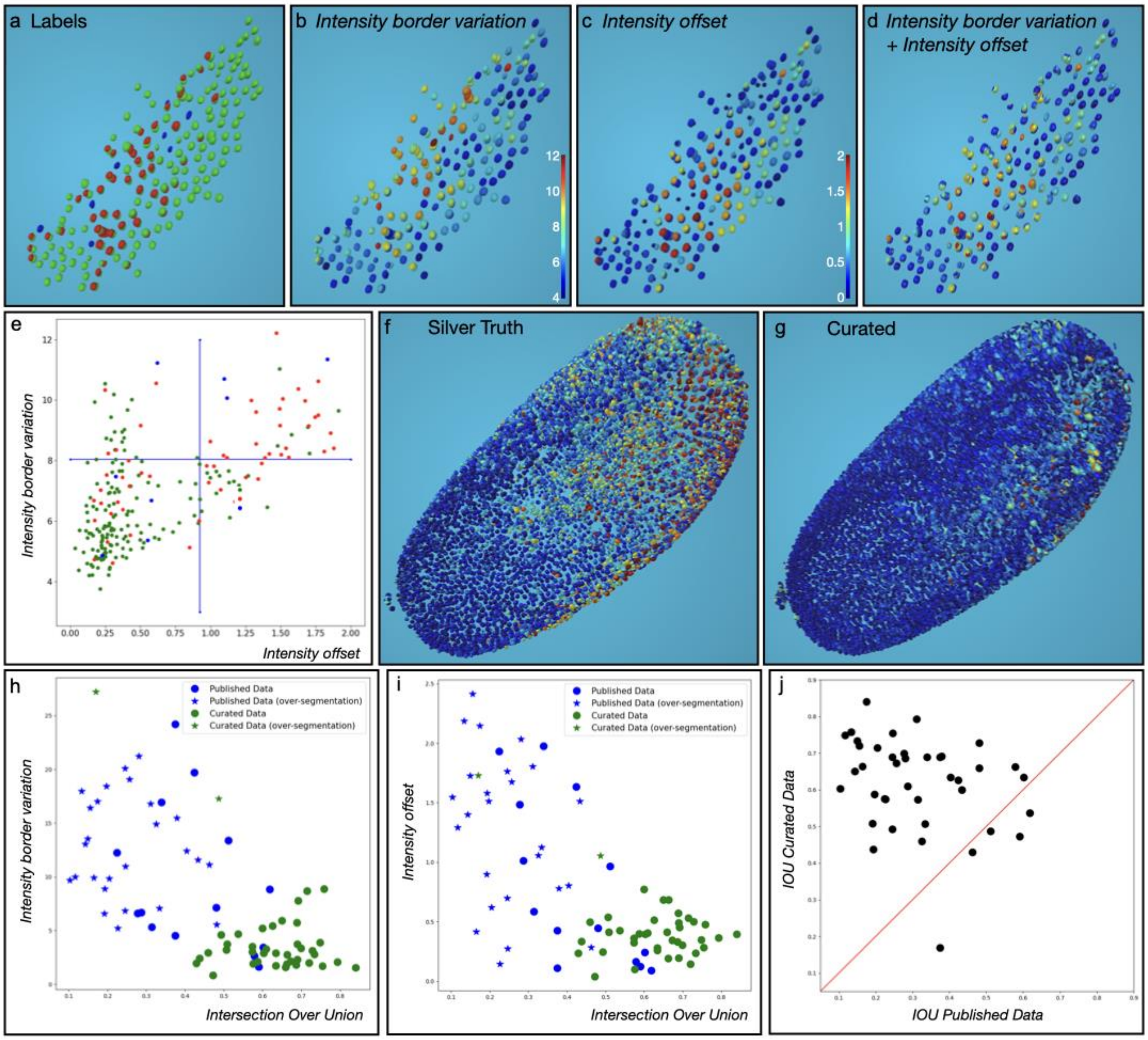


**Supplementary Figure 3: Analysis of 2 automatically computed intensity properties.**

a. Published data, Silver truth(ST) segmentation corresponding to the ROI of the Gold truth (GT). Color code : green: exact matching between a segmentation in the ST and the GT ; red: identified over-segmentation in the ST close to the GT; blue : missing cell of the GT with no correspondence in the ST.
b. Projection of the property *intensity_border_variation* onto the nuclei segmentation selected in a.
c. Projection of the property *intensity_offset* onto the same nuclei segmentation.

d. Double projection of both property *intensity_offset* and *intensity_border_variation.* Colormap same as in b. and c.
e. 2D plot of the values of both properties (*intensity_offset in X-axis and intensity_border_variation in Y-axis)* of the same selected nuclei segmentation. Colorcode is equivalent as in a. The blue line in each axis represents the last quartile of the distribution.
f. Double projection of both properties (same as in d.) on the whole ST embryo.
g. Same as in f. For the curated data obtained using the 5 steps pipeline (see Fig. 3).
h. 2D plot of 40 ground truth cells and their corresponding segmentations in both the published and curated datasets, showing *intensity_offset* property on the X-axis and Intersection over Union (IoU) on the Y-axis.
*i.* Same as in h with the *intensity_border_variation* property on the X-axis.
j. Comparison of IoU values for 40 ground truth cells and their corresponding segmentations in the published and curated datasets. Points above the red line correspond to cells with improved IoU values in the curated dataset.

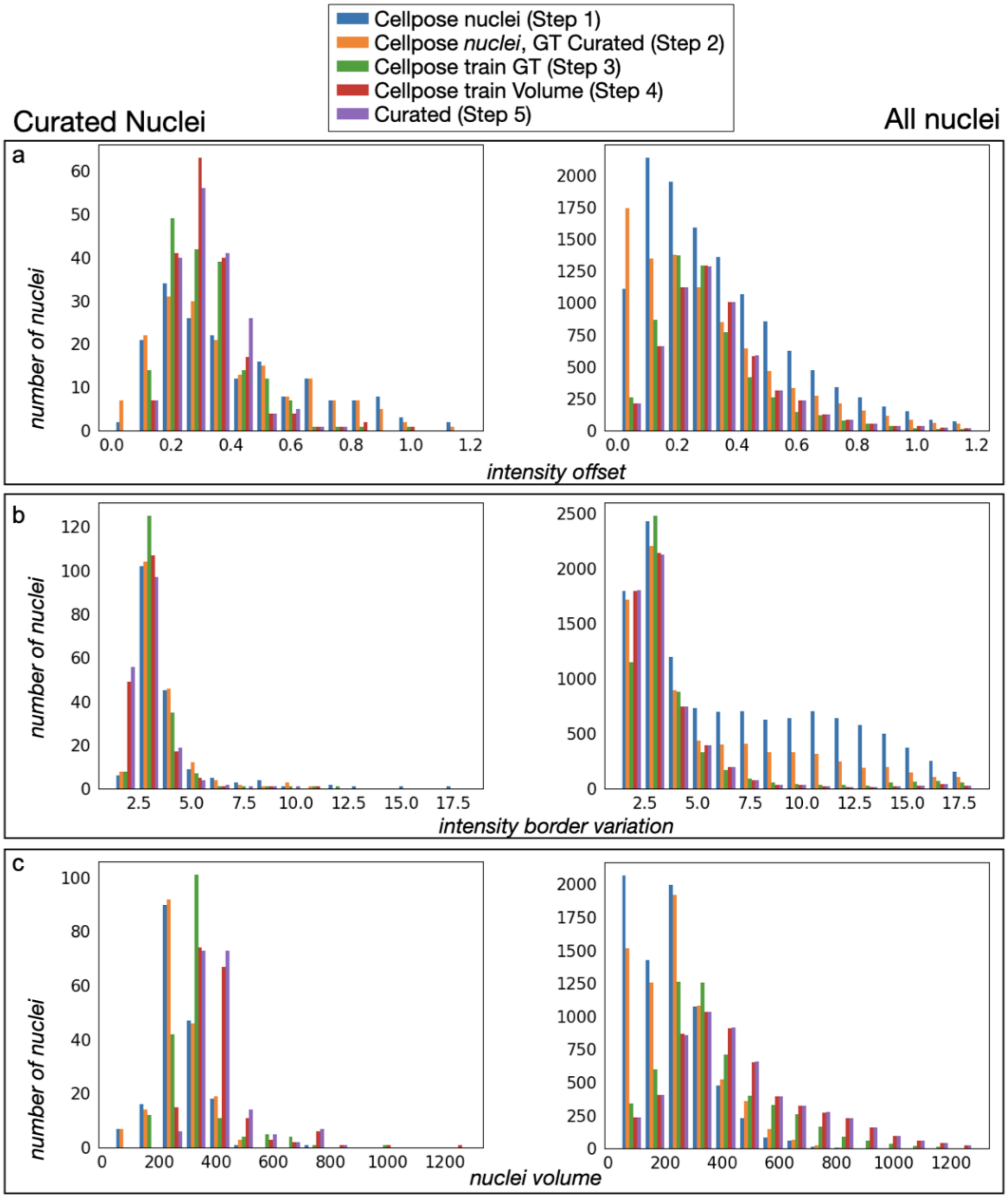


**Supplementary Figure 4: Distribution of 3 properties for the Tribolium reconstruction.**

Comparative histogram of a specific property distribution between the ST, the Step 1 and the Step 5 for the 181 curated nuclei (left) and the whole image (right).

a. Comparative histogram of the distribution of the *intensity offset* property

b. Comparative histogram of the distribution of the *intensity border deviation* property.
c. Comparative histogram of the *nuclei volume* distribution.

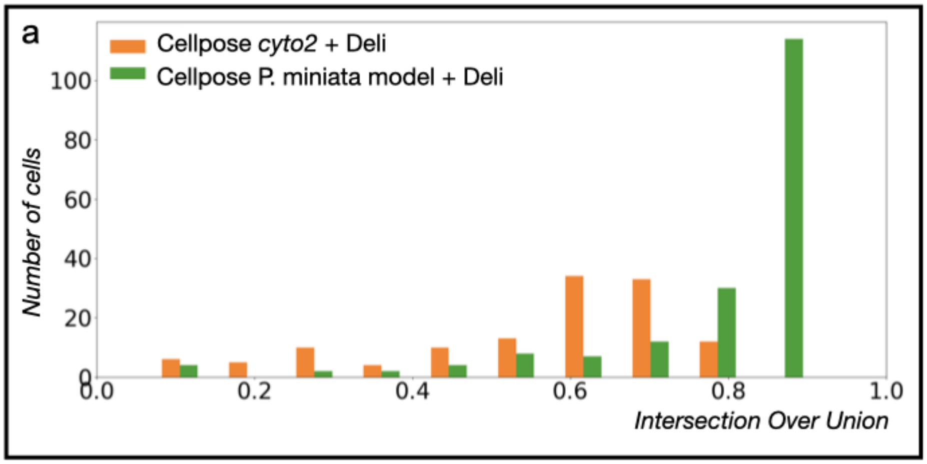


**Supplementary Figure 5: Evaluation of Segmentation in *Patiria miniata* starfish embryo.**

a. Histogram of Intersection over Union (IoU) values comparing the published dataset (gray) with the *CellPose cyto2* + *Deli* pipeline (orange) and the CellPose P. miniata model + Deli pipeline (green).

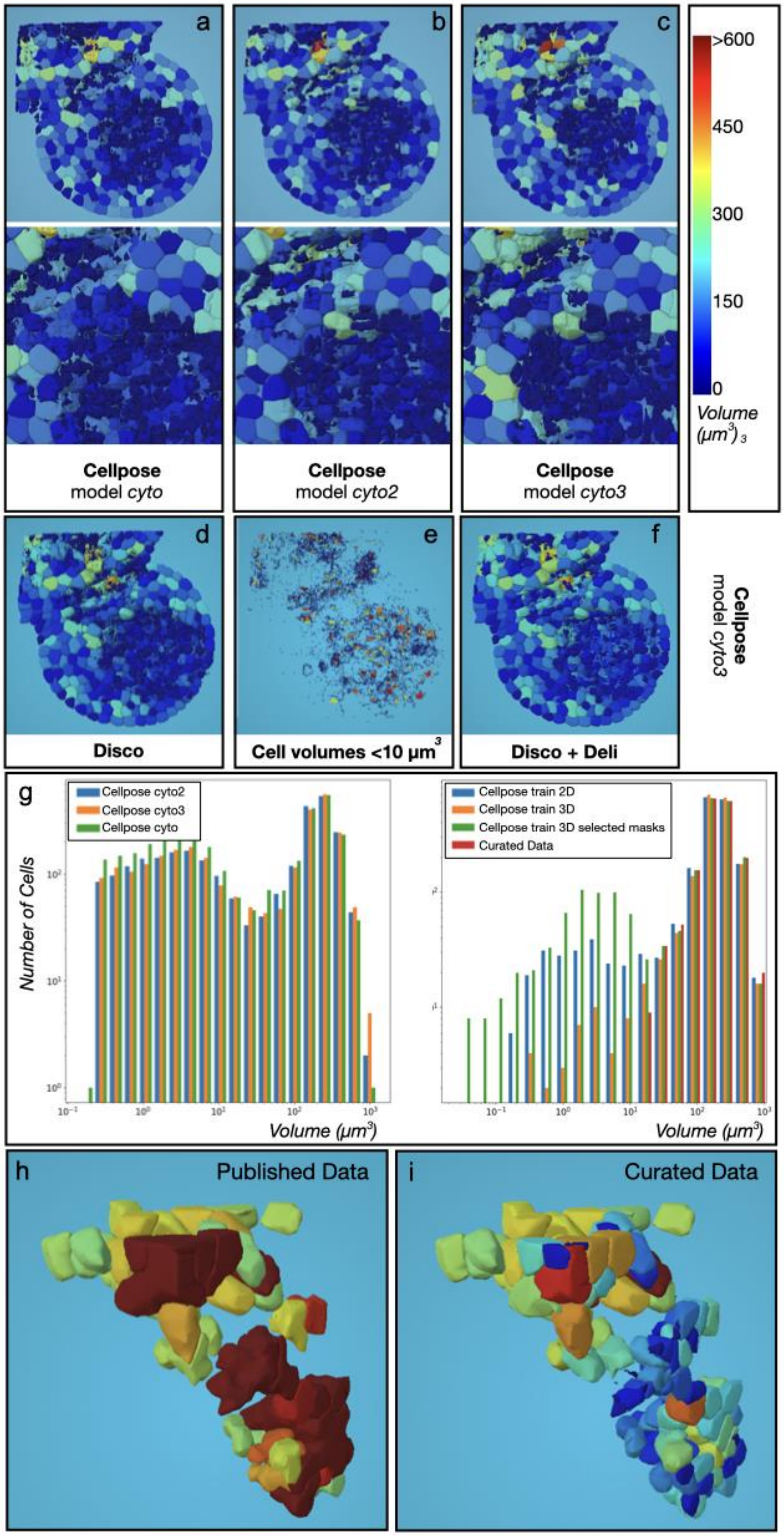
a
b
c
>600
450
300
150
0
Volume
(µm³)
Cellpose
model cyto
Cellpose
model cyto2
Cellpose
model cyto3
d
e
f
Cellpose
model cyto3
Disco
Cell volumes <10 µm³
Disco + Deli
g
Cellpose cyto2
Cellpose cyto3
Cellpose cyto
Number of Cells
Volume (µm³)
Cellpose train 2D
Cellpose train 3D
Cellpose train 3D selected masks
Curated Data
Volume (µm³)
h
Published Data
i
Curated Data

**Supplementary Figure 6: Comparison of the Cellpose models for the reconstruction shoot apical meristem of *Arabidopsis thaliana*.**

Result of the Cellpose prediction on the full image using the model *cyto(a.), cyto2(b.), cyto3(c.)*. Colorbar on the right represents the cell volume in $\mu m^3$ .

d. Result of the Cellpose prediction using the model *cyto3* with the option disconnect non connected component activated.
e. Identical as d. with the visualisation for cells with a volume <10 $\mu m^3$ .
f. Result of the Cellpose prediction using the model *cyto3* with the option disconnect non connected component activated and the option to reassign objects under a volume <10 $\mu m^3$.
g. Distribution of the segmented cell volume($\mu m^3$) for the time step 19. Left : comparison of the 3 default models of Cellpose (cyto,cyto2 and cyto3). *Cellpose train 3D*: the model was trained using XY,YZ and XZ planes and the prediction is then applied on the full 3D image. *Cellpose train 3D* selected masks: the model was trained using XY,YZ and XZ planes and the prediction is then applied only on the selected masks with volumes > 300 $\mu m^3$.
h. 3D view of the masks larger than 300 $\mu m^3$ in the published dataset.Same orientation as in Fig. 5a.
i. Same view of the corresponding masks after curation

**Supplementary Figure 7: *Manual Validation of Curation for the Caenorhabditis elegans Embryo***

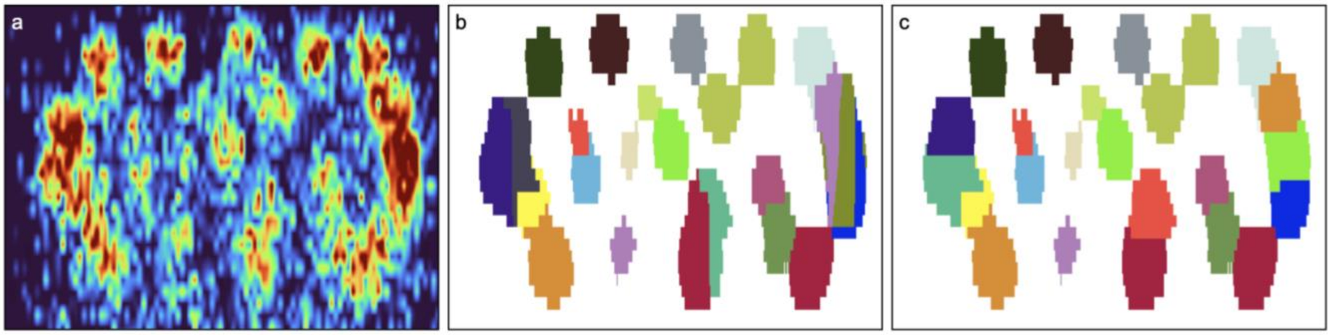


a. 2D XZ view at Y = 306 of the intensity image at time point t = 150 using Napari.
b. Same view showing the published segmentation, where vertically elongated segmentation artifacts are visually apparent.
c. Same view after curation, where these artifacts have been corrected.

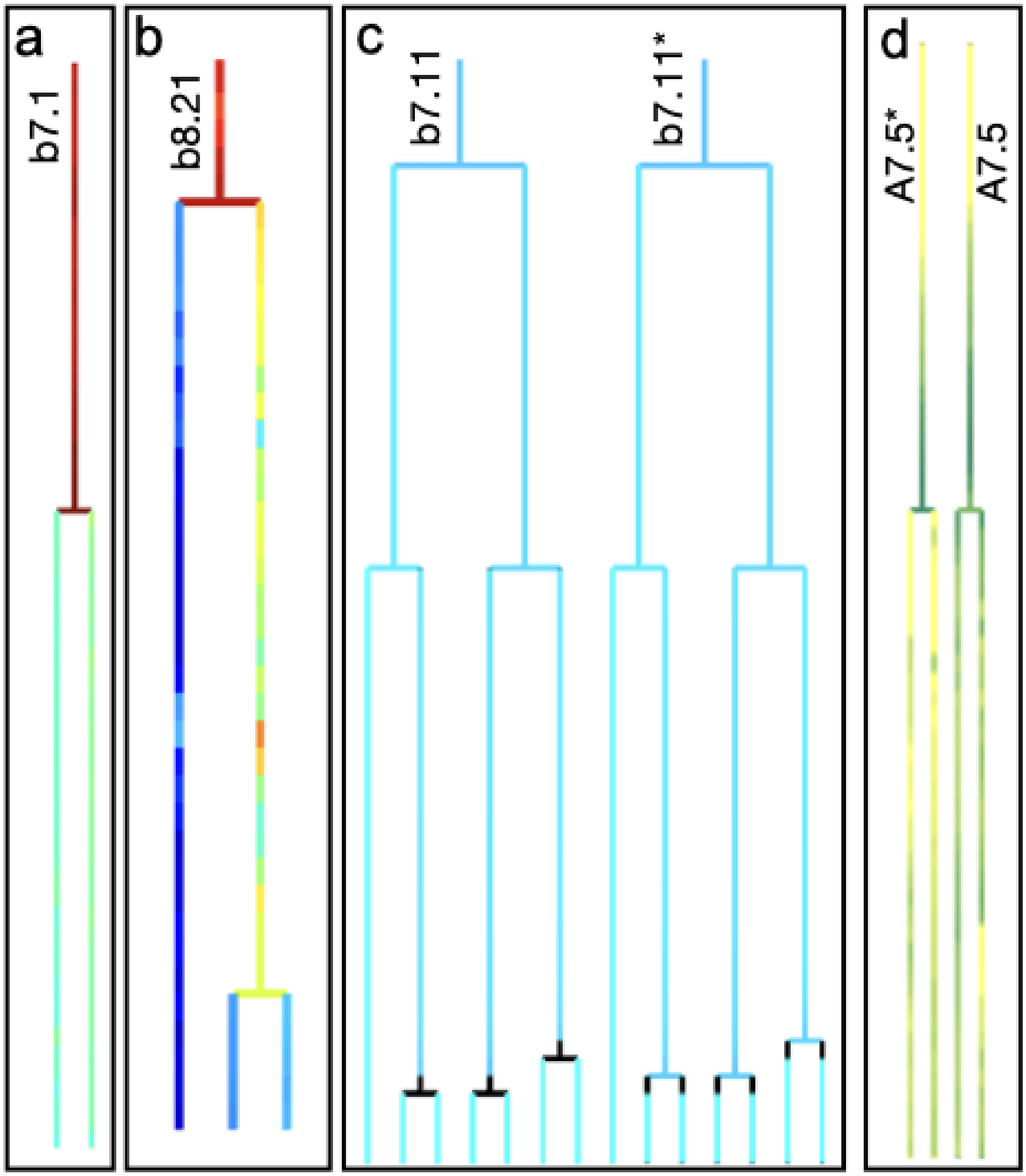


**Supplementary Figure 8: Cell lineage following curation of a *Phallusia mammillata* embryo.**

a. Cell lineage of the cell b7.1 after curation. Projection of the volume property as in Fig 7a.

b. Cell Lineage of the cell b8.21 after curation. Colormap represents the cell volume in $\mu m^3$ as in Fig 7.d
c. Cell lineage of the bilateral cells b7.11 and b7.11* after curation. Colorbar shows the lineage distance between the bilateral symmetrical cells as in Fig 7e.
d. Cell lineage of the bilateral cells a7.5 after curation. Colorbar shows the compactness property as in Fig 7f.

#### Tools Comparison

MorphoNet (MN) is a platform specifically designed for end users, requiring no programming skills. Its standalone version is a portable, code-free application, similar in spirit to Fiji (FJ).

MN provides interactive visualization of segmentations and supports both segmentation creation (via a range of plugins) and curation through a broad set of image processing tools. Unlike FJ and Napari (NP), which typically operate at the voxel level, MN performs curation exclusively at the object level. This object-centric approach enables faster and more intuitive editing of complex 3D segmentations.

MN also supports cell tracking and lineage visualization. It includes plugins for both automatic lineage generation and manual curation, similar to FJ's TrackMate plugin and NP's btrack (the latter requiring some coding knowledge).

MN offers built-in tools for visualizing and exporting morphological properties of segmented objects. In contrast to FJ, MN can directly map these properties onto segmentations using color-coded overlays, and can also project them onto lineage trees in its dedicated viewer.

Deep learning-based tools like Cellpose and StarDist are natively embedded in MN's standalone application, supporting both inference and model training with no additional installation. In comparison, FJ includes only a StarDist plugin, while NP supports both tools but typically requires command-line installation and environment setup.

Finally, all MN plugins are bundled and maintained by the development team, ensuring integration and stability. In contrast, plugin ecosystems in FJ and NP are open, allowing users to freely develop, install, and share new tools with the community.

#### Uses Cases

#### UC 1: *Tribolium castaneum* embryo cell nuclei segmentation

Curated pipeline: Due to the high number of errors in the published ST, we decided to generate *de novo* nucleus segmentation using MorphoNet. We defined a pipeline in five steps, which required only a few manual currations. Step 1- We started by launching the *Cellpose prediction* plugin using the pretrained standard *nuclei* model. Step 2- The nuclei of this segmentation corresponding to the published GT (181 nuclei) were curated with various plugins. Eight segmentations with wrong shape were recomputed using the *Binarize* plugin. One missing nucleus was added with the *BinCha* plugin which creates a new segmentation from the selected GT using a binary threshold on intensity image. Twelve segmentation errors (which were wrong segmentation boundaries between adjacent nuclei, and over-segmentations) could be corrected by using combinations of the *Fuse*, *Gaumi* and *Delete* plugins. Also, using the intensity properties, we visually identified 248 over-segmented nuclei outside of the GT, that were easily corrected with the *Fuse* plugin. Step 3- We developed a *Cellpose Train* plugin with a user-friendly interface, which we used to extend pretrained Cellpose models (here *nuclei*) using XY, XZ and YZ plane information. We added an option to train a model on selected objects within a region of interest of the image. We extended the *nuclei* model by an additional training step restricted to the 181 curated GT cells and used this new model to predict a new segmentation on the entire image. Step 4- Finally, based on this new segmentation, we automatically select nuclei with a volume between 1 000 and 10 000 voxels and perform a new Cellpose train using this selection only. Step 5- To achieve a perfect shape for each GT associated nuclei, we identified 35 nuclei with non-convex shapes using the *convexity* property, which we then refined using the *Shape Transform* plugin family. (See Supp. Mat. Plugin list).

Manual segmentation protocol: To sample a representative set of segmentation quality levels, we used the histograms of the intensity_border_variation and intensity_offset properties computed from the original Cell Tracking Challenge dataset. For each metric, one cell was randomly selected from each of the 20 histogram bins, yielding a total of 40 cells. For each selected cell, a dilated bounding box was extracted from the original intensity image, and manual ground truth segmentation was performed using Napari’s label editing tool. To detect over-segmentation, any segmented object other than the matched one that overlapped the manual ground truth by at least 5% of its volume was classified as an over-segmentation.

#### UC 2 : Whole membrane segmentation of a *Patiria miniata* starfish embryo

Original workflow: The published workflow started with the construction of a 3D Voronoi diagram based on cell centroids calculated from the nuclei channel acquisition, which was used to train a 3D ResU-Net[4] on 14 consecutive stacks from a single embryo at the 256-cell stage. The predictions obtained with this trained model on wild-type and mutant embryos from the 128- to 512-cell stages were used as input for PlantSeg[5], a watershed-based algorithm. Segmentation errors were corrected using custom Matlab scripts and the resulting segmentations were used to train a second 3D ResU-Net model, which was applied to the whole dataset. In compressed embryos, the authors choose to segment only 196 cells from a 512-cell stage embryo (Fig. 4b), the signal intensity in the embryos' center and external circumferences being considered too low for precise and objective segmentation.

Training Protocol for MorphoNet segmentation with the advanced Cellpose plugin: To train CellPose 3D on the 300 *Patiria miniata* starfish embryos kindly provided by the authors of the paper[2], we split it into 3 sub groups: a training set corresponding to 80% of the database, a testing set (10%) and a validation set (10%). We then split 3D images into 2D slides in each direction (X,Y,Z). Cellpose has to preload data in memory to convert them in the native format before training. Due to inherent memory limitation, we had to split the training process into several trials. Each trial contains 5000 random images of the training set. We run 50 trials with each of them containing 50 epochs. We started the first trial using the *cyto2* model and then each successive trial was continued to train based on the model obtained in the previous one. We then evaluate each trial on the testing set and keep the one which give the best accurate segmentation.

#### UC 4: *Caenorhabditis elegans* embryo cell nuclei segmentation

Original workflow: The dataset contains 3D stacks of intensity images and two types of ground truth annotations. A silver truth (ST) automated segmentation of each nucleus using StarryNite[11]. A gold truth (GT) manual annotation of the position of each nucleus over time, whose tracking was manually corrected with AceTree[12].

#### UC 5: *Phallusia mammillata* embryo whole-cell segmentation

Error correction: The 41 labels (which should be cells) consisting of more than one connected component were disconnected using the *Disco* plugin. Visual inspection using the *volume* property of these objects identified 7 potential cells. The others, considered as small artifacts, were removed by fusion to the cell neighbor they share most surface with using the *Deli* plugin. This plugin requires the threshold value for the minimum cell volume, which can be manually found by specifically highlighting all small cells (Fig. 7b).

Division timing issues typically arise from over-segmentation, under-segmentation or missed cells. Over Segmentations were easily corrected using the *Fuse* plugin (Fig. 7g). Using the *Propagate segmentation* family plugins, under-segmentations were corrected by backwards propagation from the first accurate segmentation of the two sister cells in the dataset up to the precise moment of division of their mother (Fig. 7h). Missing cells can be difficult to correct when signal intensity quality is poor. One correction option is to add a missing seed with the seed generator plugins (See Supp. Mat.). The expert can then apply one of several local seeded watershed algorithms to generate the missing cell segmentations (Fig. 7i). When the imaging quality is too low for this strategy to succeed, we reasoned that the bilaterally symmetrical cell may be a good mirror-image approximation of the missing cell. We thus used the *Copy-Paste* plugin to copy a cell from her symmetric and to paste it, after appropriate rotation and scaling, where a cell is missing (Fig. 7j).

Using a combination of using 264 MorphoNet plugin actions, 185 errors were corrected in 8 hours actions, leading to a better estimation of the natural variability in cell division timing (Fig.7k)

**Supplementary Movies**

For each dataset in the use cases, there is a short movie describing the curation process or parts of the curation process. All videos are available here : https://seafile.lirmm.fr/published/morphonet_data/MOVIES

#### Supplementary Movie 1: *Tribolium castaneum*

The movie shows how to train a Cellpose model using a curated sub-part of a dataset, and how to fine-tune models with successive training on specific nuclei using image properties

#### Supplementary Movie 2: *Patiria miniata*

The movie shows how to create a dataset, how to use an already existing custom Cellpose model and how to easily correct usual Cellpose segmentation errors.

##### **Supplementary Movie 3:** *Arabidopsis thaliana*

The movie shows how to train a Cellpose model on several time steps of a 3D+t dataset, and how to use it to predict under-segmentations on a specific part of a dataset.

##### **Supplementary Movie 4:** *Caenorhabditis elegans*

The movie shows how to use the lineage and image properties to easily detect segmentation issues, and how to fix them in batches for a fast curation of dense datasets.

##### **Supplementary Movie 5:** Phallusia mammillata

This movie shows how to identify and fix several issues on a segmented dataset, using the lineage viewer and a large array of plugins. It shows how to fix large curation errors using a couple of actions only.

#### Cellpose Plugins

##### Cellpose Train plugin

The current version of Cellpose has limitations for 3D model training. It requires command line or Python API usage, which is inaccessible for most experimental biologists. Additionally, Cellpose only trains on XY planes of 3D datasets, making it inefficient for anisotropic datasets with lower axial resolution, like the Arabidopsis dataset, resulting in over-segmented cells. These issues were addressed by developing a MorphoNet Cellpose Train plugin with a user-friendly interface that leverages information from the XY, XZ, and YZ planes, improving segmentation by fine-tuning pretrained models using 3D data. We added an option to make the image isotropic (by lineage interpolation) before launching the training which increases the model accuracy. Users have the possibility to train their own model on several time steps. Finally, in order to be able to use partial annotation, this plugin can be run only on selected masks such as in the Triboliutm dataset.

#### Cellpose Predict plugin

The Cellpose Predict plugin in MorphoNet is dedicated to the prediction of 3D segmented images. We implemented in MorphoNet an option to run each algorithm only on selected masks only (e.g. on cells containing segmented errors). Thus this plugin can be run only on a subpart of the dataset (such as in Arabidopsis dataset). Cellpose generates multiple unconnected components with the same label which can be removed using the Disconnected Components option (Identical as the Disco Plugin). Applying the *cyto3* model on the Arabidopsis dataset generates 13293 objects with disconnected components (Supp. Fig. 5d and e). We also add the possibility to automatically remove these small artifacts generated by Cellpose using an option to to fuse or delete the objects below a given size (It will remove these objects if they are small artefacts in the background) (Supp. Fig. 5f). This targeted approach is fastest, gives the best results and has the additional advantage of preserving the accurate cell segmentations.

### Properties

The majority of the region properties are computed from the scikit-image python package [as described here](). Properties marked with a (v) are computed on the original image, whereas others are computed on an image re-scaled by its voxel size, to give appropriate physical measurements.

- **volume** (v): Corresponds to the *area* property of scikit-image. Area of the region, i.e. number of voxels in the region.
- **volume-real**: Volume property but scaled by the voxel-size to give a real physical volume
- **volume-bbox**: Corresponds to the *area_bbox* property of scikit-image. Area of the bounding box i.e. number of voxels of bounding box scaled by voxel-area.
- **volume-filled**: Corresponds to the *area_filled* property of scikit-image. Area of the region with all holes filled in.
- **axis-major-length**: The length of the major axis of the ellipse that has the same normalized second central moments as the region.
- **axis-minor-length**: The length of the minor axis of the ellipse that has the same normalized second central moments as the region.
- **axis-ratio**: Ratio of the longest axis over the smallest axis of the label (see *axis-major-length* and *axis-minor-length*).
- **diameter** (v): The mean diameter of the region. It is the mean of axis-major-length and axis-minor-length. IMPORTANT: this value is exprimed in voxels, not physical size, so it can be of use for plugins that use voxel measurements, such as Cellpose for instance.

- **equivalent-diameter-area**: The diameter of a circle with the same area as the region.
- **euler-number**: Euler characteristic of the set of non-zero pixels. Computed as the number of connected components plus the number of holes, subtracted by the number of tunnels.
- **extent**: Ratio of pixels in the region to pixels in the total bounding box. Computed as *volume* divided by amount of rows multiplicated by amount of columns in bounding box
- **connected-neighbors**: number of connected other labels.
- **convexity**: distance to convexity. Computed as volume of the convex hull (smallest convex polygon that encloses the region) divided by the volume of the region.
- **roughness**: mean of the absolute values of the region minus the closing (dilation followed by erosion) of the region.
- **compactness**: computed as $\frac{region\ surface\ area^3}{36 \times \pi * volume^2}$
- **smoothness**: computed as $\frac{region\ surface\ area}{volume^{\frac{2}{3}}}$
- **Intensity-max** (v): Value with the greatest intensity in the region .
- **Intensity-mean** (v): Value with the mean intensity in the region.
- **Intensity-min** (v): Value with the least intensity in the region.
- **Intensity-border-variation** (v): the standard deviation of the intensity images only at the border of the segmentation.
- **Intensity-offset** (v): the euclidean distance between the gravity center of the intensity images and the geometrical center of the segmentation.
- **Lineage-distance** : Requires a lineage property, as well as the *cell_name* property, containing the Conklin[14] naming of cells. Compute the tree-edit distance between lineage trees[13] of symmetrical cells.
- **Bbox** (v): bounding box of the region (tuple).

#### Curation

The MorphoNet standalone application allows users to perform the 3D images curation using a new paradigm. The concept is based on a duality between the segmented image and the construction of 3D mesh objects for each individual label. While the meshes object are extremely powerful for 3D visualization and interaction, the 3D segmented image remains the most standard data type for images labels. The curation is performed in 4 steps:

1. Users identify their issues to curate using the viewer based on the interaction with the messages (using Lineage or Intensity Images)
2. Users identify and launch the most appropriate plugin to solve their issue
3. The plugin perform the curation directly inside the segmented image
4. MorphoNet automatically recompute the modification of the meshes of the labeled which have been modify by the plugin and finally refresh the window

Each plugin performs a modification of the 3D segmented images. The modification is done locally on the backup of the segmented images stored inside the MorphoNet application, not on the original data. Finally, users can export their curation as new segmented images. Meshes can also be exported in standard formats including OBJ, STL, and PLY.

MorphoNet includes a list of various 3D plugins described below. We also add the possibility for any bio-image analysis to simply develop, test and add their own plugins (See Supplementary Table 1 ). The python environment already contains several libraries such as numpy[15], scikit-image[16], scikit-learn[17], tensorflow[18] and pytorch[19].

##### Curation Plugins

MorphoNet already included a list of various 3D plugins which are classified by family types: *De Novo Segmentation, De Novo Seed, Segmentation from Seeds, Segmentation Correction, Propagate Segmentation, Edit Temporal Links and Shape Transform.*

**De Novo Segmentation :** these plugins function without any previous segmentation and directly create a segmentation from the 3D intensity images. Alternatively, they can override any segmented image (or a part of) when used on already segmented images (Full online documentation).

***Stardist-Predict:** Perform nuclei segmentation on intensity images*

This plugin uses intensity images of the nucleus from a local dataset, to compute segmentations of the nucleus, using the 3D Stardist deep learning algorithm[20]. The default demo model of stardist can be used, as well as custom models (which can be created by the Stardist-Train plugin ).

***Stardist-Train:** Train the Stardist model for nuclei on your own data*

This plugin allows users to train their own Stardist model of their 3D datasets. Using the 3D intensity image(s) with the corresponding segmentation(s), users can train their own models. The models can subsequently be used in the Stardist-Predict plugin to perform nuclei segmentation prediction on 3d intensity images[20].

***CellPose-Predict:** Perform membrane segmentation on intensity images*

This plugin uses an intensity image of the membranes from a local dataset at a specific time point, to compute a segmentation of the membranes, using the 3D CellPose deep learning algorithm[8]. Users can apply CellPose on a 3D selected Mask to apply it to a part of a segmentation. By default, the plugin also disconnects all non-connex labels in the generated segmentation, and deletes all segmentations below a certain volume (in voxels), the same way it does in the *Disco* and *Deli* plugins. Each of these operations can be disabled with the plugin parameters.

***CellPose-Train:** Perform membrane segmentation on intensity images*

This plugin allows users to train their own model (from the models provided by CellPose) on their own 3D datasets. With an intensity image and the corresponding segmentation, users can re-train a Cellpose model to obtain their own model, trained on their images. The model will be trained by using 2D images, which are the image stacks on the XY, XZ and YZ planes of the 3D images. Users can apply the plugin on a 3D selected Mask to train on a sub-part of a segmentation.

The model you output from this plugin can then be used in the Cellpose-Predict plugin, by inputting it in the pretrained_model parameter.

**Mars:** Perform a seeded watershed segmentation on intensity images

This plugin uses an intensity image from a local dataset at a specific time point, to perform a segmentation using a seeded watershed algorithm[21].

**Binarize**: Apply a threshold to intensity image

This plugin applies a threshold to the intensity image and then creates labels on each connected component above the threshold. This binarization can be applied on a given mask to run it on a sub-part of the image.

**BinCha**: Apply a binary threshold on the other channel from selected objects

This plugin creates a new segmentation from each mask of the selected objects using a binary threshold on intensity image in the desired channel. Alternatively, you can also do the thresholding on the centroid of each object, with a bounding box of input radius.

**BinBox**:  Binarize intensity images and label each object inside a bounding box

This plugin applies a threshold to intensity image on a bounding box and creates new labels on each connected component above the threshold.

***De Novo* Seed:** these plugins automatically create seeds (e.q. 3D points within the 3D images), which can be then used by the plugins in the Segmentation from seeds family (Full  online documentation).

***Seedio:*** *Create seeds from minimum local intensity images on the selected objects*

This plugin generates seeds that can be used in other plugins (mainly watershed segmentation). Users have to select objects (or label them) to generate seeds. Seeds are generated at the minima inside the selected object.

***Seedin:*** *Create seeds from minimum local intensity images  (without selected objects)*

This plugin generates seeds that can be used in other plugins (mainly watershed segmentation). Seeds are generated at the minimum intensity where no segmentation labels are found (in the background)

***Seedis:*** *Create seeds from the maximum distance to the border of the selected objects (without intensity images)*

This plugin generates seeds that can be used in other plugins (mainly watershed segmentation). It computes the distance to the border of the selected objects and then extracts the maxima. N seeds are generated at the maximal distance inside the selected object (N being the number of seeds to generate), if the distance (between seeds) is above the threshold given by the min_distance parameter (in voxels, not physical size).

***Seedax:*** *Create Seeds on the long axis of the selected objects (without intensity images)*

This plugin generates seeds that can be used in other plugins (mainly watershed segmentation). The longest axis of the segmentation shape is computed, and then split in N segments (N being the number of seeds in parameter). Seeds are generated at the contact points of the segments. It requires the user to select or label objects on MorphoNet.

***Seedero:*** *Create seeds from the erosion of selected objects (without intensity images)*

This plugin generates seeds that can be used in other (mainly segmentation) algorithms. This plugin applies multiple erosion steps of each selected object, until objects can be separated into multiple unconnected parts. Then a seed is placed at the barycenter of each individual sub-part of the segmentation.

#### Segmentation from Seeds: these plugins can generate segmentations using seeds (mostly with a watershed algorithm[22]). Seeds can be either added by a plugin or manually on the interface ([Full online documentation]).

***Watio:*** *Perform a watershed segmentation on intensity images on selected objects*

This plugin creates new objects using a watershed algorithm from seed generated using a plugin or manually placed in the MorphoNet Viewer inside selected objects.

The watershed algorithm generates new objects based on the intensity image and replaces the selected objects. If the new generated objects are under the volume threshold defined by the user, the object is not created.

***Wati:*** *Perform a watershed segmentation on intensity images (without selected objects)*

This plugin creates new objects using a watershed algorithm from seed(s) generated or placed in the MorphoNet Viewer. The watershed algorithm generates new objects using the intensity image for each seed that is in the background. If the new generated objects are under a volume threshold defined by the user, the object is not created.

***Wato:*** *Perform a watershed segmentation on selected objects (without intensity images)*

This plugin creates new objects using a watershed algorithm from seed(s) generated using a plugin or placed in the MorphoNet Viewer inside selected objects. The watershed algorithm generates new objects using the segmentation image for each seed and replaces the selected objects. If the new generated objects are under the volume threshold defined by the user, the object is not created.

***Wata:*** *Perform a watershed segmentation (without intensity images and without selected objects)*

This plugin creates new objects using a watershed algorithm from seed(s) generated using a plugin or placed in the MorphoNet Viewer. The watershed algorithm generates new objects using the segmentation image for each seed inside a box that is not in another object. If the new generated objects are under the volume threshold defined by the user, the object is not created.

**Segmentation Correction:** a various list of plugins to perform actions at the level of the selected objects (fusion, deletion, split, copy-paste, etc.) (Full online documentation).

***Fuse:*** *Fuse the selected objects*

This plugin performs the fusion of selected objects into a single one. Multiple selected objects will be fused together at the current time point. If objects are labeled it will apply a fusion between all objects sharing the same label (for each individual time points)

***Gaumi:*** *Split the selected objects using probability distribution*

This plugin calculates the gaussian mixture model probability distribution on selected objects in order to split them into several objects which will replace the selected ones.

***Disco:*** *Split the selected objects for unconnected objects*

This plugin can be used to split any object made up of several unconnected sub-objects.

***Splax:*** *Split the selected objects in the middle of a given axis*

This plugin splits any selected objects into two new objects in the middle of one of the given image axes.

***Delete:*** *Delete the selected objects*

This plugin removes any selected objects from the segmented images. Users can select objects at the current time point, or label any objects to delete at several time points. The background value (usually 0) will replace the object voxels inside the segmented image.

***Deli:*** *Delete any objects below a certain size*

This plugin removes all objects that are under a certain volume (in voxel count) from the segmented images.

***Copy-Paste:*** *Copy a selected object and apply a transformation to the copy*

This plugin gives the possibility to copy an object (a segmented cell for example) and paste it at another time step and/or another location. The object can be moved, rotated and scaled on the MorphoNet interface.

***Match:*** *Matching objects across multiple channels*

This plugin allows you to match the elements of the same object across several channels. It will give the selected objects a matching label in the segmented image. Can be used in batch by labeling objects together with label groups.

**Shape Transform:** these plugins allow the user to change the shape of objects, with Morphological operators (Erode, Dilate, etc.) and even manually (Full online documentation).

***Dilate:** Dilate the selected objects*

This plugin performs the dilation morphological operator on each individual selected object

***Erode:** Erode the selected objects*

This plugin performs the erosion morphological operator on each individual selected object

***Open:** Open the selected objects*

This plugin performs the opening morphological operator on each individual selected object

***Close:** Close the selected objects*

This plugin performs the closing morphological operator on each individual selected object

***Convex:** Make selected objects convex*

This plugin computes the convex voxel hull of each individual selected object

***Deform:** Apply a manual deformation on the selected object*

This plugin allows the user to manually deform a selected object using mesh deformation. This plugin must be used with the mesh morphing menu and its various tools. The mesh morphing menu allows the user to manually deform a selected object of your dataset by applying various transformations (move vertices, extrude, hollow, ...) with the mouse pointer. Once the user is satisfied with the deformation, this plugin computes the transformation(s) applied to the mesh to the segmented image in the dataset, and regenerates the mesh object using this segmented data.

**Propagate Segmentation:** these plugins allow the user to propagate a good segmentation at a given time to other time points (Full online documentation).

***Propa:** Propagate selected eroded objects through time on the next object (with or without intensity images)*

This plugin propagates labeled objects at the current time point through time. It requires applying a specific label to the objects to propagate at the current time (named the source objects) And then label the corresponding objects on which to propagate at the next or previous time points (named the destination objects). The plugin can be executed forward (source objects are at the beginning of the time range) or backward (source objects are at the end of the time range). The source objects are eroded until they fit into destination objects at the next time point, and then a watershed is computed. The watershed algorithm can be computed using an intensity image by checking the "use intensity" box. The new objects created by a watershed will replace the destination objects

***Prope:*** *Propagate selected eroded objects through time on the next empty area with intensity images)*

This plugin propagates labeled objects at the current time point through time, filling empty space. The selected object(s) (named the source objects) will be propagated Forward in time or Backward in time by choosing the appropriate time_direction parameter. The source objects are then copied to the segmentation of the next/previous time (the intersection with the rest of the segmentation is removed), and finally a watershed algorithm is applied, using the corresponding intensity image. New objects are created in the background of the segmentation.

#### Edit Temporal Links: these plugins can create or modify temporal information in order for example to curate a cell lineage tree (Full online documentation).

***Addlink:*** *Create temporal links between labeled objects*

This plugin creates temporal links at several time points on objects sharing the same label. After the execution, the lineage property is updated.

***Delink:*** *Delete temporal links on labeled objects*

This plugin delete temporal links at several time points on objects sharing the same label. After the execution, the lineage property is updated.

***Tracko:*** *Create temporal links using the overlap between all objects*

This plugin creates a complete object lineage using the maximum of overlap between objects. The overlap is calculated between the bounding box enveloping each object. After the execution, the lineage property is updated.

##### Create a new plugin

It is possible for any user to develop their own curation plugins in python, and integrate them in their version of the MorphoNet standalone application. Users that wish to develop their own plugins can do so by following a tutorial on the MorphoNet website help page, on the link below.

https://morphonet.org/help_api?menu=morphonetplot#plugins

#### Data availability

All the original datasets and the curations presented are available here:
https://seafile.lirmm.fr/published/morphonet_data/DATASETS

#### Code availability

The MorphoNet platform is composed of several Open Source repositories:

- The frontend 3D Viewer code and Unity3D project is available here : https://gitlab.inria.fr/MorphoNet/morphonet_unity
- The python API code, which contains the python backend of the standalone application is available here : https://gitlab.inria.fr/MorphoNet/morphonet_api
- The lineage viewer code and Unity3D project is available here: https://gitlab.inria.fr/MorphoNet/morphonet_lineage
